# Infection-Induced Elevation of Gut Glycosaminoglycans Fosters Microbiota Expansion in *Drosophila melanogaster*

**DOI:** 10.1101/2025.06.24.661374

**Authors:** Aranzazu Arias-Rojas, Marko Rubinić, Jasmin Albiez, Dagmar Frahm, Robert Hurwitz, Igor Iatsenko

## Abstract

While host genetics influence the composition of intestinal microbial communities, host genetic factors controlling the abundance of intestinal commensals remain to be determined. Here, we performed a genome-wide association (GWA) study in the fruit fly *Drosophila melanogaster* to identify host genetic variants linked to the abundance of *Lactiplantibacillus plantarum* – a major gut commensal of fruit flies. Our GWA study uncovered significant association between polymorphisms in genes involved in heparan sulfate synthesis and *L. plantarum* load. RNAi mediated knockdown of some of these genes resulted in reduced heparan sulfate synthesis and *L. plantarum* abundance. Mechanistically, heparan sulfate facilitates adhesion of *L. plantarum* to host epithelium and promotes biofilm formation. We further showed that infection induces heparan sulfate synthesis by the host via activation of the Nf-kB immune signaling cascade. Increased availability of heparan sulfate during infection results in the expansion of *L. plantarum* population in the gut and protection of the host from intestinal pathogens via colonization resistance. Furthermore, heparan sulfate is required for infection-induced expression of immune effectors and for prevention of intestinal dysplasia. These findings underscore heparan sulfate as a crucial modulator of intestinal homeostasis, pivotal in microbiota control, intestinal defense, and epithelial renewal.

## Introduction

Host epithelial surfaces are abundantly colonized by microbial communities called microbiota. Microbiota is an integral part of the organism’s physiology affecting immune, metabolic, behavioral, neurological, and disease traits among others ^1–3^. Environmental factors, like diet and xenobiotics, and ecological interactions between microbes shape substantially the composition of microbiota ^4,5^. Considering the benefits provided by some microbiota members, selection for host genotypes that favor a beneficial microbiome should also be possible. Indeed, population-based studies demonstrated the role of genetic component in determining gut microbiota composition, and a proportion of bacterial taxa is heritable ^6,7^. Genome-wide association studies in humans identified dozens of loci associated with microbiome composition and the abundance of specific taxa ^8–10^. Genes with a role in immune regulation and barrier defense were enriched among the identified host factors associated with microbiome variation ^11^. However, given little reproducibility across studies, besides well- replicated associations with the *LCT* and *ABO* genes ^12^, we know little about host genetic factors that control the abundance of specific gut microbes. Considering that an excessive increase in abundance of particular gut bacteria has been associated with intestinal disease, immunodeficiency, and negative outcome of antibiotic treatments ^13–15^, identification of host genes and mechanisms behind microbiota control is of paramount importance for prevention of microbiota overgrowth and its consequences.

Genome-wide association studies are a promising approach to find the associations between host genes and the microbiome. However, in humans, such studies require large sample size and are complicated by the fact that environmental factors, such as diet, can strongly and rapidly alter gut community composition ^12,16^, making it challenging to differentiate the effect of environment and host genetics. Model organisms, like the fruit fly *Drosophila melanogaster*, can help to overcome these challenges. First, GWA studies in *Drosophila* can be done with a small sample size thanks to a community tool called *Drosophila* Genetic Reference Panel (DGRP). DGRP is a collection of 205 genetically-heterogenous lines with sequenced genomes widely used to perform genotype-phenotype associations ^17,18^. Second, fruit flies have a low-diversity gut microbiota, of <30 taxa and commonly dominated by five species in the genera *Acetobacter* and *Lactobacillus*, which are readily culturable ^19–21^. Studies of host-microbiota interactions in *Drosophila* are also facilitated by the simplicity of generating germ-free and gnotobiotic animals ^22,23^. Hence, precise control of the environment and interactions between microbes allows the dissection of the specific associations between host genetics and the microbiome in fruit fly model. GWA approach identified several *Drosophila* genes linked to the abundance of *Acetobacter tropicalis* ^24^. However, since *A. tropicalis* was used along with another 4 species, it is not known how the presence of the other microbes influenced the results. Early at el ^25^ used gnotobiotic DGRP lines colonized with a single bacterial strain that is known to reside in the fly gut and found that genes strongly associated with bacterial levels are largely involved in neuronal function, neuronal morphogenesis and development. Considering the small sample size, many associations likely were not detected. Hence, the full potential of *Drosophila* GWAS to identify the associations between host genes and the microbiome remains to be exploited.

Notably, with the exception of antimicrobial peptide Defensin potentially associated with *A. tropicalis* control ^24^, none of the other GWAS-identified genetic variants were linked to immune response or processes known to control bacterial load in flies. These processes include acidic pH and digestive enzymes that create an inhospitable environment to many bacteria ^26–28^, peritrophic matrix-a chitinous barrier, that prevents microbial access to the epithelium ^29^, and clearance of pathogens via intestinal peristalsis ^30–33^. Production of reactive oxygen species (ROS) ^34,35^, antimicrobial peptides (AMPs) ^36^, and iron-sequestration proteins ^37,38^ in specific gut regions represent inducible immune reactions controlling microbial proliferation. Disruptions of many of these defensive barriers were linked to uncontrolled growth of gut bacteria, dysbiosis, and short lifespan. For instance, flies lacking AMPs are short-lived due to microbiome dysbiosis ^39,40^. Similarly, experimental conditions that induce high densities of specific gut bacteria in *Drosophila* can impair gut function and depress fly life span ^41^. For example, *Gluconobacter morbifer* in flies with an overactive immune system ^42^ and a monoassociation of immunocompromised mutants with *Lactiplantibacillus plantarum* bacteria that induces production of ROS ^43^. Thus, the abundance of specific microbes is a crucial factor in host-microbe interactions that needs to be controlled to maintain homeostasis.

Studying microbial factors implicated in host colonization can also be a fruitful approach to identify host determinants of microbial control. For instance, Gutiérrez-García et al. identified glycan-binding adhesins used by *L. plantarum* to colonize the symbiotic niche in the foregut of *Drosophila* ^44,45^. While exact identity of host glycans bound by *L. plantarum* remains to be determined, they should contain *N*-acetylglucosamine, as blocking *N*-acetylglucosamine by feeding flies with lectins inhibited gut colonization by *L. plantarum.* This work together with the other studies ^46,47^ on prominent role of glycosaminoglycans (GAGs) in host-microbe interactions suggest that host control of GAG availability/composition might be a strategy to control the diversity and abundance of gut microbiome.

Resistance to host AMPs is another mechanism that allows gut commensals to colonize the gut ^48,49^. Previously, we characterized several *L. plantarum* mutants sensitive to AMPs and showed that the abundance of these mutants was severely reduced in infected but not in control uninfected flies. Notably, wild-type *L. plantarum* even benefited from infection and increased significantly in quantity ^48,50^. Hence, the host genetic architecture underlying the abundance of microbiota is likely to be different under infection conditions.

In this study, we uncovered host genetic determinants of *L. plantarum* abundance under homeostatic and infection conditions, revealing a previously unappreciated role of heparan sulfate in microbiota control, host defense, and epithelial renewal.

## Results

### *Drosophila* genotypes vary in the load of commensals

Under natural conditions the abundance of a particular commensal in the gut is determined by the interplay of three main factors: host genotype, environment, and interactions between microbiome community members. To determine the effect of host genotype only and exclude the contribution of the other two factors, we maintained a controlled environment and introduced a single commensal strain into the gut, thus eliminating the interactions with other community members. To test whether fly genotype affects the quantity of intestinal commensal bacteria, we generated 92 gnotobiotic DGRP lines colonized by a single commensal strain – *L. plantarum* WJL (*Lp*). Each *Lp*-colonized DGRP line was exposed to sucrose (basal condition, referred to as unchallenged, ‘UC’) or infected with fly natural pathogen *Pectobacterium carotovorum* (*Ecc15)* for 6 h (Figure 1a), allowing us to test the effect of host genotype on commensal load under homeostatic and infection conditions. As shown in Figure S1a and S1b, DGRP lines showed variation in *Lp* load under basal (UC) and *Ecc15*-infected conditions. Interestingly, some DGRP lines (line 28) with high *Lp* load under UC conditions had low *Lp* load after infection and another way around. This suggests that *Lp* abundance after infection can’t be predicted from UC conditions, indicating distinct genetic architecture determining commensal load after infection. Consistent with our observation in wild-type flies ^48^, we found that most of DGRP lines exhibited an increase in *Lp* load after infection relative to basal conditions (Figure 1b).

**Figure 1.**
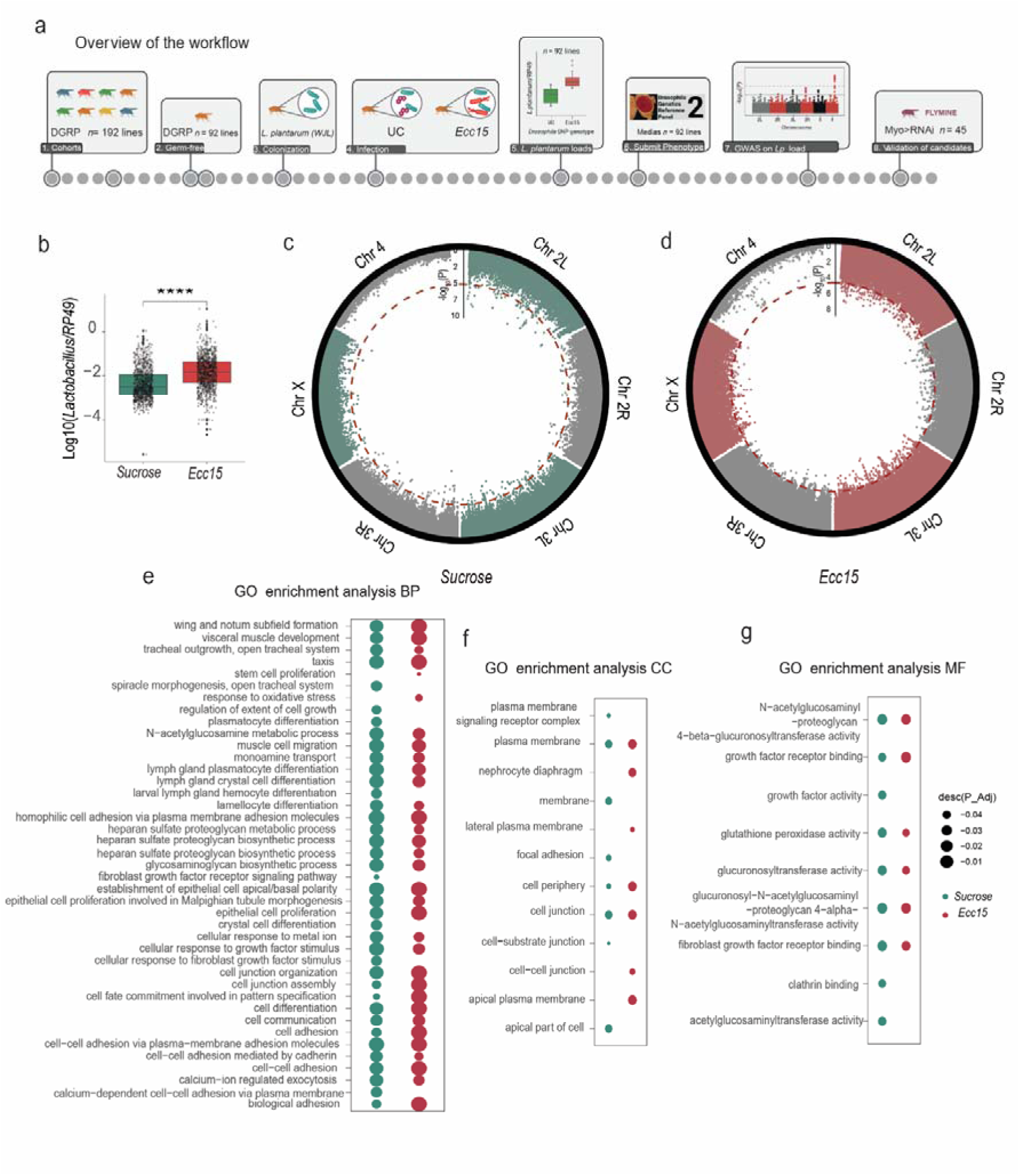
Genome-wide association study identified host genetic variants controlling *L. plantarum* abundance. (**a**) Overview of the GWAS workflow. (**b**) Levels of *L. plantarum* among the DGRP lines after sucrose and infection with *Ecc15* (n=8 per line). (**c** and **d**) Circular Manhattan plots show the hits per chromosome (1-4, and X), the red dotted lines show the cut-off on the p-value of -log10(5), and the dots above the line show the top-tier genes. (**e-g**) Gene ontology (GO) enrichment analysis of candidate genes associated with *L. plantarum* levels at the Biological Process (e), Cell Component (f), and Molecular Function (g) levels. Circles show the p-adjusted value significance, and the color shows the cluster of the term per treatment.

### Identification of candidate genes

The DGRP facilitates rapid GWA analyses using a quantitative phenotype via submission of a data set to the online webtool ^17^. To determine polymorphisms in the DGRP population that affect abundance of *Lp*, we submitted log_10_-transformed *Lp* titres for analysis. We tested a total of 1861607 polymorphisms in the sucrose and *Ecc15* infection analysis. We employed a genome-wide suggestive *P*-value threshold of 10CC, commonly used in DGRP studies, as it balances false positive control with the ability to detect potential associations in moderate-sized datasets. Using this *P*-value, we obtained a total of 343 associated polymorphisms for basal and 1000 associated polymorphisms for *Ecc15* condition from the GWA analyses (Table S1, S2, Figure 1c, 1d).

Of the 343 polymorphisms associated to the sucrose condition, 139 variants are intergenic (40.52%), 163 are within introns 47.5%; 3 are within exons (0.87%); 5 are within the 5C UTR (1.45%); and 9 are within the 3C UTR (2.6%). For *Ecc15* infection, 736 variants are intergenic (73.6%), 962 are within introns (96.2%); 12 are within exons (1.2%); 46 are within the 5C UTR (4.6%); and 76 are within the 3C UTR (7.6%) To further dissect the genetic architecture underlying *Lp* abundance, we performed a Gene Ontology (GO) analysis using the top associated variants. Using Ensembl annotations, we assigned each variant to a single gene. We analyzed sucrose (156) and *Ecc15* conditions (409) separately. With FlyMine ^51^, we found that several GO categories are enriched in each gene set (Figure 1e-g). Vast majority of the GO categories are shared between the two datasets. However, GO terms “response to oxidative stress” and “stem cell proliferation” were specifically enriched in *Ecc15* gene set, suggesting that these processes might be important for *Lp* load control during infection. GO terms related to hemocyte differentiation were specifically enriched in sucrose gene set. Notably, many GO categories showing significant fold enrichment are involved in heparan sulfate and N-acetylglucosamine metabolism, and general cellular growth and development.

Next, we selected several top candidate genes from different GO categories and tested their role in the control of *Lp* abundance under basal and infection conditions. For this purpose, we generated *Lp*-colonized gnotobiotic RNAi lines with gut-specific candidate gene knock down and assessed *Lp* load after sucrose and *Ecc15* infection relative to control RNAi (Figure S2a). We could validate the role of several tested candidate genes in the control of *Lp*. For example, lines Dip-theta, Liprin alpha, Naked, Prosap, Luna, nAChRalpha6 and Syt4 exhibited reduced *Lp* load compared to control under both sucrose and *Ecc15* infection (Figure S2b). Other genes, like *Dip-epsilon* are important for *Lp* load control only during infection as they showed the same *Lp* abundance as control under basal conditions (Figure S2b). We could also find genes (*cnc, CG11373, Glut1*) that control *Lp* levels under basal conditions but not during infection (Figure S2b). Hence, we could validate several of GWA-determined candidates and identify new host genetic determinants of commensal load under basal and infection conditions.

### Heparan sulfate mediates gut colonization by *Lp* via enhancing adhesion and biofilm formation

Since our GO analysis identified significant enrichment of genes related to heparan sulfate metabolism and considering the prominent role of heparan sulfate pathway in mediating host- microbe interactions ^47,52^, we decided to mechanistically dissect its contribution to the control of *Lp* load in flies. As a starting point, we assessed *Lp* abundance in *ttv (tout-velu)* RNAi. Ttv was identified as a candidate from GWA and encodes a glycosyltransferase required for the biosynthesis of heparan sulfate by the addition of beta-1-4-linked glucuronic acid (GlcA) and alpha-1-4-linked N-acetylglucosamine (GlcNAc) units to nascent heparan sulfate chains ^53^. Consistent with a previous report ^54^, we could not detect heparan sulfate in *ttv* RNAi line with antibodies against heparan sulfate (Figure S3a), demonstrating that *ttv* knockdown efficiently inhibits heparan sulfate synthesis in the gut. *Ttv* RNAi showed reduced *Lp* abundance relative to control under both basal and infection conditions, suggesting that Ttv is required for *Lp* colonization and infection-induced boost in commensal load (Figure 2a). Additionally, we estimated *Lp* abundance in other RNAi lines with knock down of several genes in heparan sulfate synthesis pathway. Most of these lines behaved similar to *ttv* and exhibited reduced *Lp* abundance, particularly after infection (Figure S3b). These results suggest that heparan sulfate pathway is involved in the control of *Lp* abundance under basal and infection conditions and genetic disruption of heparan sulfate pathway lowers *Lp* load via reducing heparan sulfate availability. To test whether heparan sulfate itself is involved in interactions with *Lp* rather than secondary consequences of pathway disruption, we pre-fed flies with heparan sulfate prior to *Lp* colonization and found that this results in elevated *Lp* abundance as compared to flies pre-fed with sucrose (Figure 2b).

**Figure 2.**
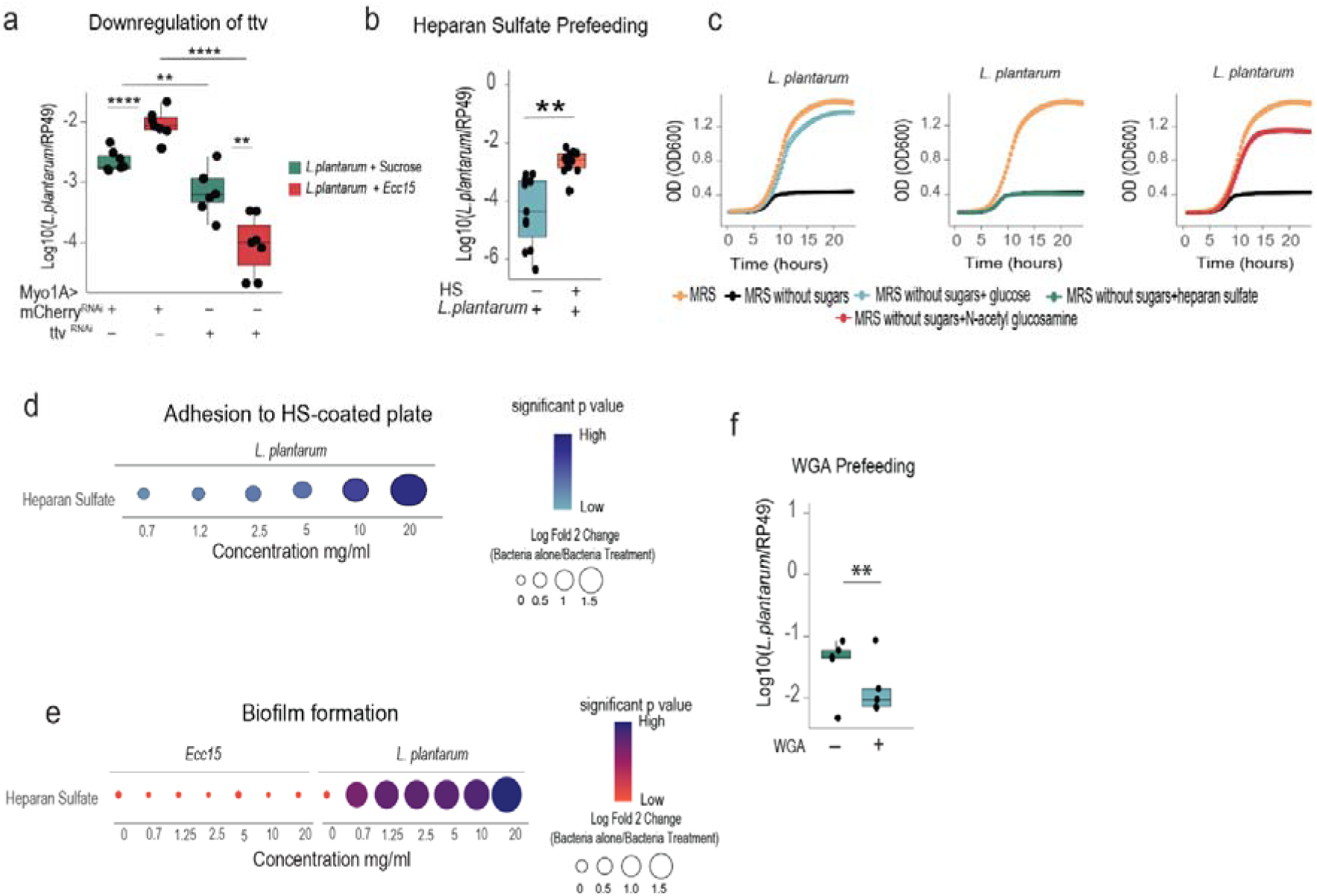
Heparan sulfate mediates gut colonization by *L. plantarum* via enhancing adhesion and biofilm formation. (**a**) *L. plantarum* loads in Myo1A>mCherry RNAi and Myo1A>ttv RNAi germ-free flies colonized with *L. plantarum* at 6 h after sucrose or *Ecc15* treatment (n=7). (**b**) *L. plantarum* loads in germ-free wild-type flies that were pre-fed with heparan sulfate or sucrose (n=12) prior to colonization with *L. plantarum*. (**c**) Growth curves of *L. plantarum* in modified MRS media supplemented with specific sugars as sole carbohydrate sources. (**d**) *L. plantarum* adhesion to plates coated with different concentrations of heparan sulfate. Fold changes in the absolute number of OD570 of crystal violet staining are shown. The color shows the significance of p-values. (**e**) Biofilm formation expressed as fold changes in the absolute number of OD570 of crystal violet staining when *L. plantarum* was cultured with different concentrations of heparan sulfate. The color shows the significance of p-values. (**f**) *L. plantarum* loads in germ-free wild-type flies that were pre-fed with wheat germ agglutinin (WGA) or sucrose (n=5) prior to colonization with *L. plantarum*.

Next, we explored how heparan sulfate could affect *Lp* abundance in the gut. First, we tested whether *Lp* could utilize heparan sulfate as a nutrient by growing bacteria in modified MRS medium containing heparan sulfate as a sole carbohydrate source. While *Lp* grew well on glucose, we could not detect any growth when heparan sulfate was used, suggesting that *Lp* can’t utilize heparan sulfate as a nutrient source (Figure 2c). Second, we tested whether *Lp* could use heparan sulfate as adhesion molecule. Indeed, we found that *Lp* exhibited dose-dependent increase in adhesion to heparan sulfate-coated wells of 96-well plate (Figure 2d). Additionally, being motivated by *Staphylococcus aureus* studies which showed that heparan sulfate increases biofilm formation ^55^, we explored whether heparan sulfate could facilitate *Lp* biofilm formation. We indeed detected an enhanced biofilm formation by *Lp* but not by *Ecc15* in the presence of heparan sulfate (Figure 2e). To further confirm the role of heparan sulfate in *Lp* adhesion, we blocked host glycans by pre-feeding flies with Wheat germ agglutinin (WGA). WGA binds primarily to *N-*acetylglucosamine, a constituent of heparan sulfate, and was shown before to inhibit gut colonization by *Lp* ^44^. We observed reduced abundance of *Lp* in flies pre-fed with WGA prior to *Lp* colonization (Figure 2f). These results support a role of heparan sulfate in *Lp* adhesion to the host epithelia.

### Infection-induced heparan sulfate synthesis contributes to the expansion of *Lp* during infection

Considering our results that *Ecc15* infection increases *Lp* load in *ttv*-dependent manner, we hypothesized that *Ecc15* might induce heparan sulfate synthesis, consequently promoting gut colonization by *Lp*. To substantiate this hypothesis, we assessed *Lp* load after heparan sulfate feeding, *Ecc15* infection, and combined treatment. Pre-feeding flies with heparan sulfate resulted in a similar to infection increased abundance of *Lp* (Figure 3a). Combined *Ecc15* and heparan sulfate treatment led to the same *<u>Lp</u>* abundance as single treatments (Figure 3a), suggesting that infection and heparan sulfate increase *Lp* abundance via the same mechanism. In further support of our hypothesis, we found the expression of *ttv* (Figure 3b) and of several other heparan sulfate pathway genes (Figure S3c) was induced by *Ecc15* in guts of axenic flies. This induced expression was particularly prominent in the guts of *Lp*-colonized flies (Figure S3c). Consistent with gene expression, quantification of heparan sulfate in the guts using Heparin Red Ultra Kit showed significantly elevated amount of heparan sulfate in the guts of *Ecc15*-infected flies as compared to axenic flies and further increase in *Lp*-colonized flies after *Ecc15* infection (Figure 3c). We further confirmed these results using immunostaining with Anti-Heparan sulfate antibody (Figure 3d). Hence, *Ecc15* increases heparan sulfate synthesis in *Drosophila* gut. To illustrate that such an increase is responsible for *Lp* expansion, we pre-treated *Lp* with heparan sulfate prior to gut colonization anticipating that such treatment would saturate heparan sulfate-binding receptors on the surface of *Lp* and reduce gut colonization. Indeed, we observed that such treatment prevented an increase in *Lp* abundance after infection (Figure 3e). Pre-feeding flies with WGA similarly abrogated *Lp* increase in infected flies (Figure 3f). These results illustrate that *Lp* binding to heparan sulfate is necessary to benefit from infection-induced synthesis of heparan sulfate.

**Figure 3.**
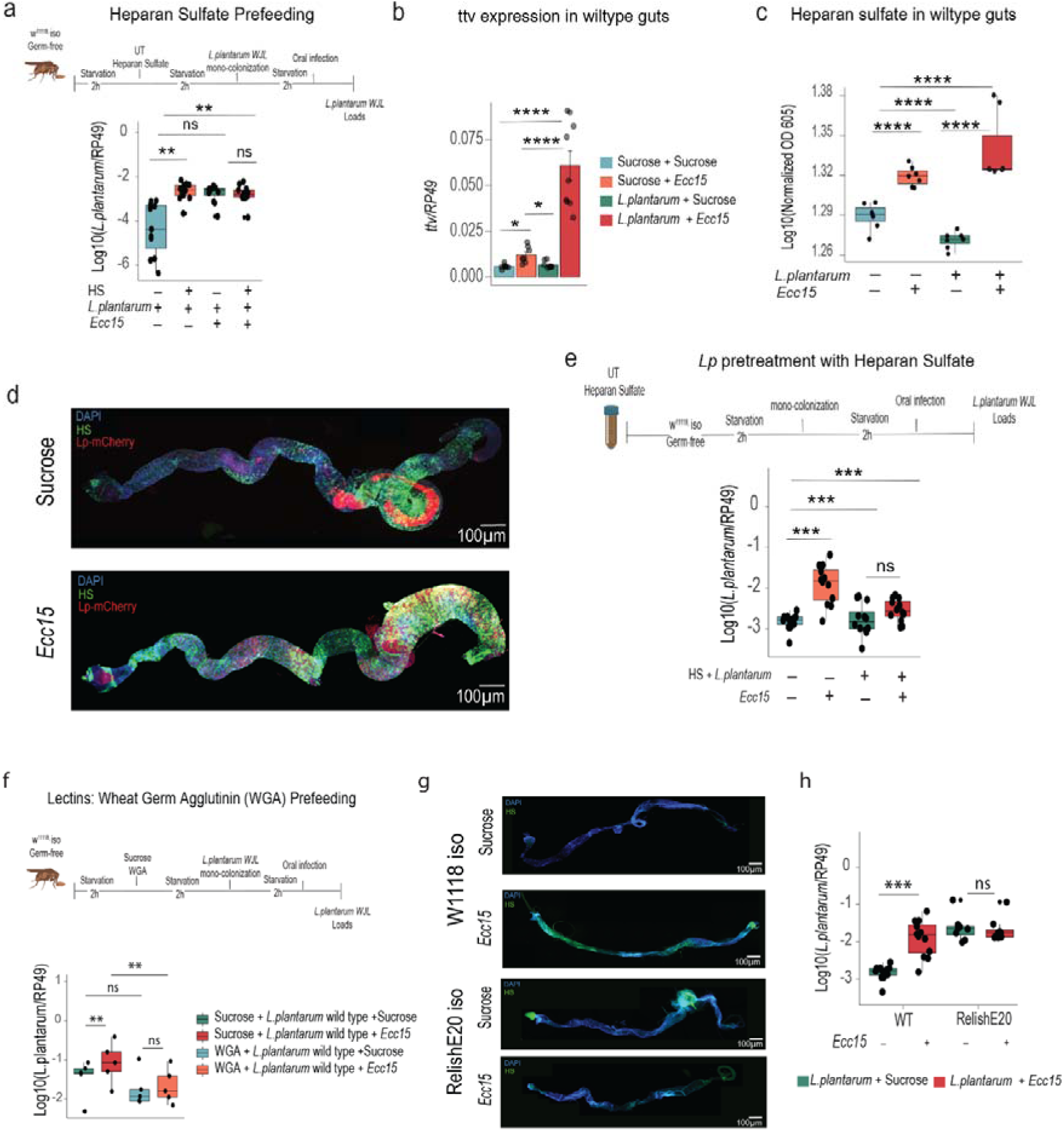
Infection-induced heparan sulfate synthesis contributes to the expansion of *L. plantarum* during infection (**a**) *L. plantarum* loads in germ-free wild-type flies that were pre-fed with heparan sulfate or sucrose (n=12) prior to colonization with *L. plantarum* and either treated with sucrose or *Ecc15* for 6h. (**b**) Expression of *ttv* gene in wild-type guts treated with sucrose only, colonized with *L. plantarum*, and treated with either sucrose again or infected with *Ecc15* and collected at 6h (n=8). (**c**) Heparan sulfate quantification with Heparin Red kit in germ-free wild-type guts colonized with *L. plantarum* or not and treated with sucrose or *Ecc16* for 6h (n=7). (**d**) Representative images of wild-type guts colonized with *L. plantarum* WJL-mCherry and treated with sucrose or *Ecc15*. Immunostaining with anti-heparan sulfate antibodies showed increased heparan sulfate signal (HS) in infected guts. (**e**) *L. plantarum* loads in wild-type germ-free flies colonized with *L. plantarum* pre-treated with heparan sulfate or MRS prior to colonization (n=12). (**f**) *L. plantarum* loads in germ-free wild-type flies that were pre-fed with wheat germ agglutinin (WGA) or sucrose prior to colonization with *L. plantarum* and either treated with sucrose or infection for 6h (n=5). (**g**) Representative images of heparan sulfate staining in guts from wild-type and *RelishE20* germ-free flies colonized with *L. plantarum* and treated with sucrose or *Ecc15* for 6h. (**h**) *L. plantarum* loads in wild-type and *RelishE20* germ-free flies colonized with *L. plantarum* and treated with sucrose or *Ecc15* for 6h (n=8).

Next, we investigated how *Ecc15* induces heparan sulfate synthesis pathway and tested whether the major immune pathway in the fly gut – immune deficiency pathway (Imd), is required for heparan sulfate induction by *Ecc15*. Flies lacking major transcription factor of Imd pathway – Relish showed abrogated expression of many heparan sulfate pathway genes that were induced in wild-type flies (Figure S3c). Immunostaining revealed only a weak signal for heparan sulfate in *Relish* mutant and no noticeable increase after infection (Figure 3g). These results illustrate that *Ecc15* induces heparan sulfate synthesis via Imd pathway. Consistent with a deficiency in infection-induced heparan sulfate synthesis, *Relish* mutant in contrast to wild-type flies failed to support *Lp* expansion after infection (Figure 3h). Overall, these results support a model where intestinal infection increases *Lp* abundance in the gut via Imd-dependent induction of heparan sulfate synthesis.

We wondered if *Ecc15,* besides promoting heparan sulfate synthesis, can make heparan sulfate more accessible for *Lp* to utilize as nutrient source. Such possibility is feasible, given that N-acetylglucosamine-a sugar constituent of heparan sulfate can be hydrolysed by *Lp* and be utilized as a sole carbon source (Fig 2c). We assumed that if *Ecc15* can digest heparan sulfate into simple sugars accessible to *Lp*, then, heparan sulfate pre-digested by *Ecc15* should be able to support *Lp* growth. However, we found no support for this assumption as *Lp* growth was not improved by *Ecc15*-treated compared to control heparan sulfate (Figure S4a). Furthermore, we used reversed-phase high-performance liquid chromatography (rp-HPLC) and derivatization with 2-aminoacridone (2-AMAC) as a method to test whether *Ecc15* can release any glycans from heparan sulfate. While we could detect derivatization product in the control sample where heparan sulfate was incubated with heparinase III, no products were detected in heparan sulfate samples treated with supernatant or pellet of *Ecc15* (Figure S4b).

Additionally, GAG analysis of control and infected flies has not identified any differences in the composition (Figure S4c), suggesting that *Ecc15* infection increases *Lp* abundance by promoting heparan sulfate synthesis rather than by digesting heparan sulfate into accessible aminosugars.

### Adhesion to heparan sulfate is necessary for *Lp* expansion in the gut during infection

To provide more direct evidence that *Lp* binding to heparan sulfate is involved in gut colonization, we decided to isolate *Lp* mutants with reduced adhesion to heparan sulfate and tested their ability to colonize flies. To identify *Lp* factors involved in adhesion to heparan sulfate, we performed a genetic screen for mutants with unaffected growth but impaired in the ability to adhere to heparan sulfate-coated plate (Figure 4a). We screened our previously- generated transposon mutant library ^48^ and identified five mutants: PL1_B10 (Redox-sensing transcription repressor), PL7_G12 (Multiple antibiotic resistance regulator), PL9_C12 (Prophage P1 protein 60, holin), PL20_D2 (extracellular transglycosylase), PL25_C9 (intergenic insertion between XynC and XpkA) (Figure 4b, Figure 4c). These mutants showed comparable to wild-type *Lp* growth (Figure S5a) but had reduced biofilm formation in the presence of heparan sulfate (Figure 4d) and reduced ability to adhere to heparan sulfate- coated plate (Figure 4e). Importantly, while under unchallenged condition these mutants reached the same abundance in the gut as wild-type *Lp*, their load in comparison with wild- type *Lp* was significantly reduced in infected flies (Figure 4f). Thus, in contrast to wild-type *Lp*, these mutants are not able to benefit from infection-induced heparan sulfate synthesis due to impaired adhesion. Notably, all five mutants exhibited similar to wild-type resistance to polymyxin B (Figure S5b), thus excluding increased susceptibility to antimicrobials as a potential reason for decreased colonization ability during infection. Hence, our screen not only identified *Lp* factors involved in adhesion to heparan sulfate but also further supported the importance of such adhesion for bacterial persistence in the gut during infection.

**Figure 4.**
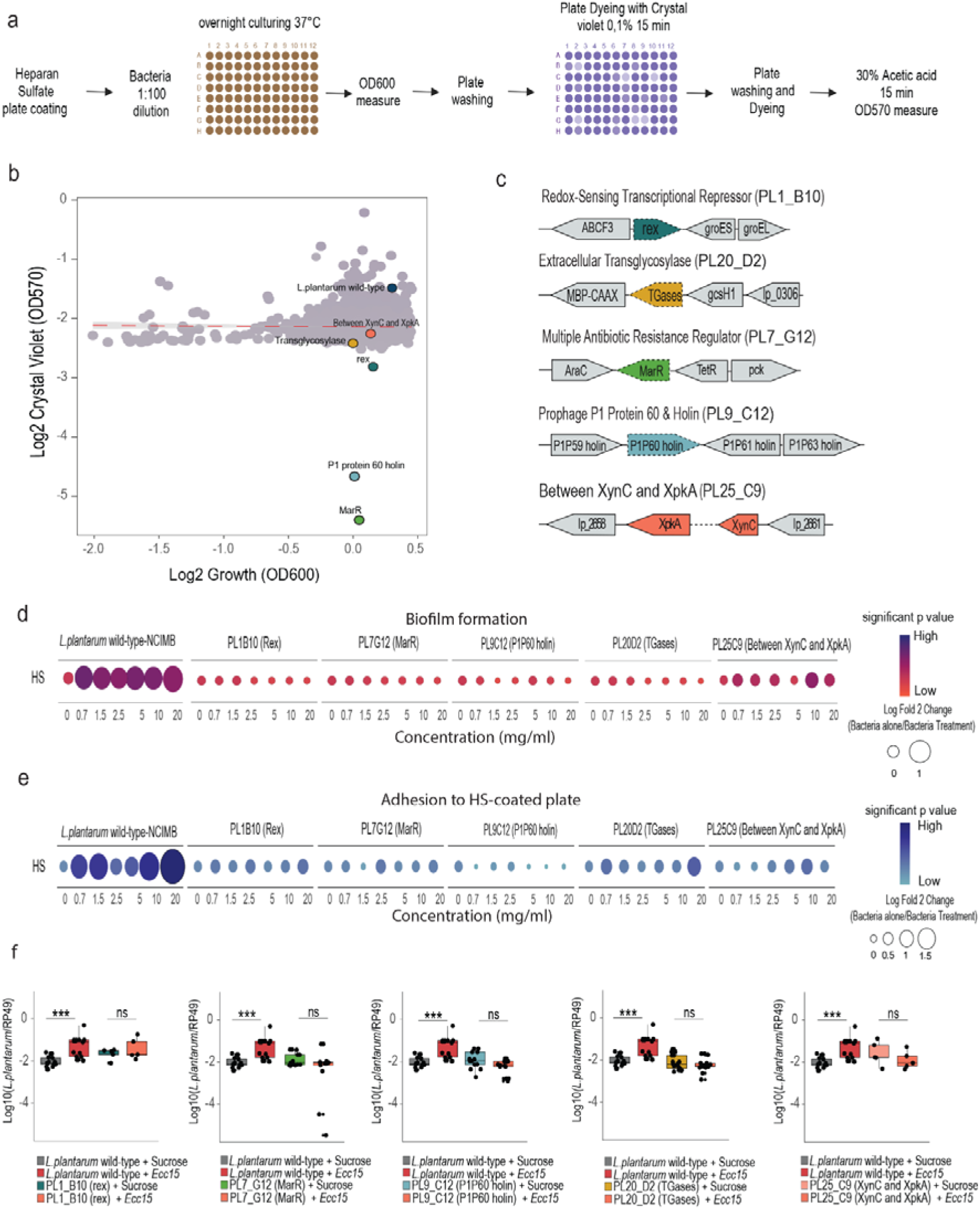
Adhesion to heparan sulfate is necessary for *L. plantarum* expansion during infection. (**a**) Overview of the screening procedure for *L. plantarum* transposon mutants with reduced adhesion to heparan sulfate-coated plates. (**b**) Scatter plot of Log-transformed relationship between bacterial growth and biofilm formation for each mutant. The red dashed line shows a LOESS regression fit. Selected mutants are labelled. (**c**) Schematic illustration of the genes and their surrounding genomic regions implicated in biofilm formation. (**d**) Biofilm formation by different *L. plantarum* transposon mutants expressed as fold changes in the absolute number of OD570 of crystal violet staining when *L. plantarum* was cultured with different concentrations of heparan sulfate. The color shows the significance of p-values. (**e**) Adhesion of *L. plantarum* transposon mutants to plates coated with different concentrations of heparan sulfate expressed as fold changes in the absolute number of OD570 of crystal violet staining. The color shows the significance of p-values. (**f**) *L. plantarum* loads in germ-free flies colonized with either *L. plantarum* wild-type or transposon mutants at 6h post treatment with sucrose or *Ecc15*.

### Heparan sulfate plays a role in intestinal defense via regulation of AMP expression and intestinal renewal

Given that heparan sulfate production is increased after infection and is controlled by the major immune response pathway in the gut – Imd, we hypothesized that heparan sulfate might play a role in host defense against intestinal infection. To test this hypothesis, we used *ttv* RNAi to reduce heparan sulfate synthesis in the gut and assessed survival of these flies to *P. entomophila* intestinal infection. *Ttv* RNAi in enterocytes (Figure 5a and S6a), enteroendocrine cells (Figure 5b), intestinal stem cells (ISCs) and progenitors (Figure 5c) led to significantly increased susceptibility of flies to *P. entomophila* intestinal infection. This effect was not specific to *ttv*, as knockdown of *botv, sotv, slf*, and *sulf* (Figure S6b-e) in enterocytes similarly resulted in enhanced susceptibility of flies to *P. entomophila* infection. Hence, heparan sulfate synthesis pathway is necessary for flies to survive intestinal infection. To obtain further insights into the underlying mechanisms, we investigated the effect of heparan sulfate synthesis pathway disruption on two key processes necessary for fly survival to intestinal infection: expression of AMPs and intestinal epithelial repair. As shown in Figure 5d, while pathogens induced AMP expression as exemplified by *Dpt* in control flies, such *Dpt* expression was abolished in flies with *ttv* knockdown in enterocytes, demonstrating that heparan sulfate synthesis pathway is necessary for AMP induction during infection. Transient burst of ISCs proliferation to repair epithelial damage is an important aspect of *Drosophila* response to infection ^36^. Using *escargot* (*esg*)*-GAL4-*driven GFP expression which marks ISCs and progenitor enteroblasts, we assessed the impact of *ttv* RNAi on stem cell proliferation under basal and infection conditions. The *esg*-*Gal4^ts^*-driven expression of *ttv* RNAi led to a significant increase in the GFP signal as compared to control RNAi already under basal conditions (Figure 5e). *Ecc15* infection expectedly triggered ISC proliferation as quantified by the increased GFP signal in control line (Figure 5e). Such increase, however, was not observed in *ttv* RNAi line (Figure 5e). Similar result was obtained with *botv* RNAi (Figure S6f). These results are consistent with previous studies ^56,57^ and suggest that disruption of heparan sulfate synthesis pathway leads to intestinal hyperplasia which was associated with increased susceptibility to gut infections ^58^.

**Figure 5.**
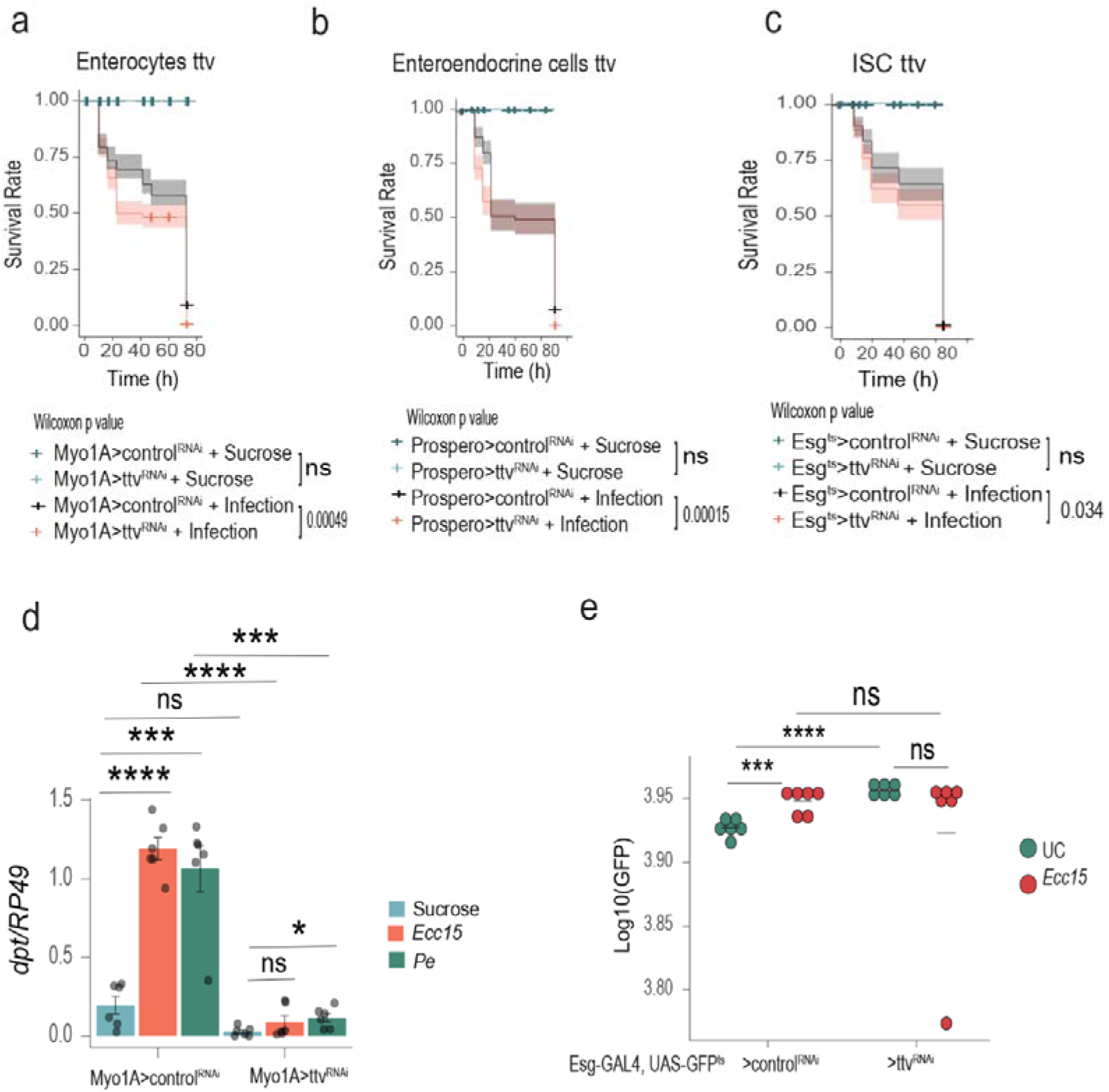
Heparan sulfate plays a role in intestinal defense via regulation of AMP expression and intestinal renewal. (**a**) Survival curves of Myo1A>attp40 RNAi and Myo1A>ttv RNAi flies exposed to either sucrose or *P. entomophila* infection (n=8). (**b**) Survival curves of Prospero>attp40 RNAi and Prospero>ttv RNAi flies exposed to either sucrose or *P. entomophila* infection (n=7). (**c**) Survival curves of Esg>attp40 RNAi and Esg>ttv RNAi flies exposed to either sucrose or *P. entomophila* infection (n=7). (**d**) *Dpt* expression in Myo1A>attp40 RNAi and Myo1A>ttv RNAi flies exposed to either sucrose, *P. entomophila,* or *Ecc15* infection for 6h (n=6). (**e**) Quantification of escargot positive cells (GFP positive cells) in Esg>attp40 RNAi and Esg>ttv RNAi treated with either sucrose or *Ecc15* for 16h (n=6).

Overall, our results show that knockdown of heparan sulfate synthesis pathway increases susceptibility of flies to intestinal infection via suppression of AMP induction and repair of epithelial damage.

## Discussion

We identified heparan sulfate as a key mediator of *Drosophila* gut colonization by a prominent commensal *Lp*. Mechanistically, *Lp* uses heparan sulfate as an adhesion molecule which also enhances biofilm formation. During infection, activation of Imd pathway leads to the increased synthesis of heparan sulfate and consequent boost of *Lp* abundance. Furthermore, heparan sulfate synthesis pathway is required for infection-induced AMP expression and for prevention of intestinal dysplasia. Hence, heparan sulfate synthesis pathway plays a dual role in *Drosophila* intestinal defense (Figure 6). First, it modulates immune response and repair pathways. Second, it promotes colonization resistance against pathogens by increasing the abundance of commensal with previously reported antagonistic activity ^26,59,60^.

**Figure 6.**
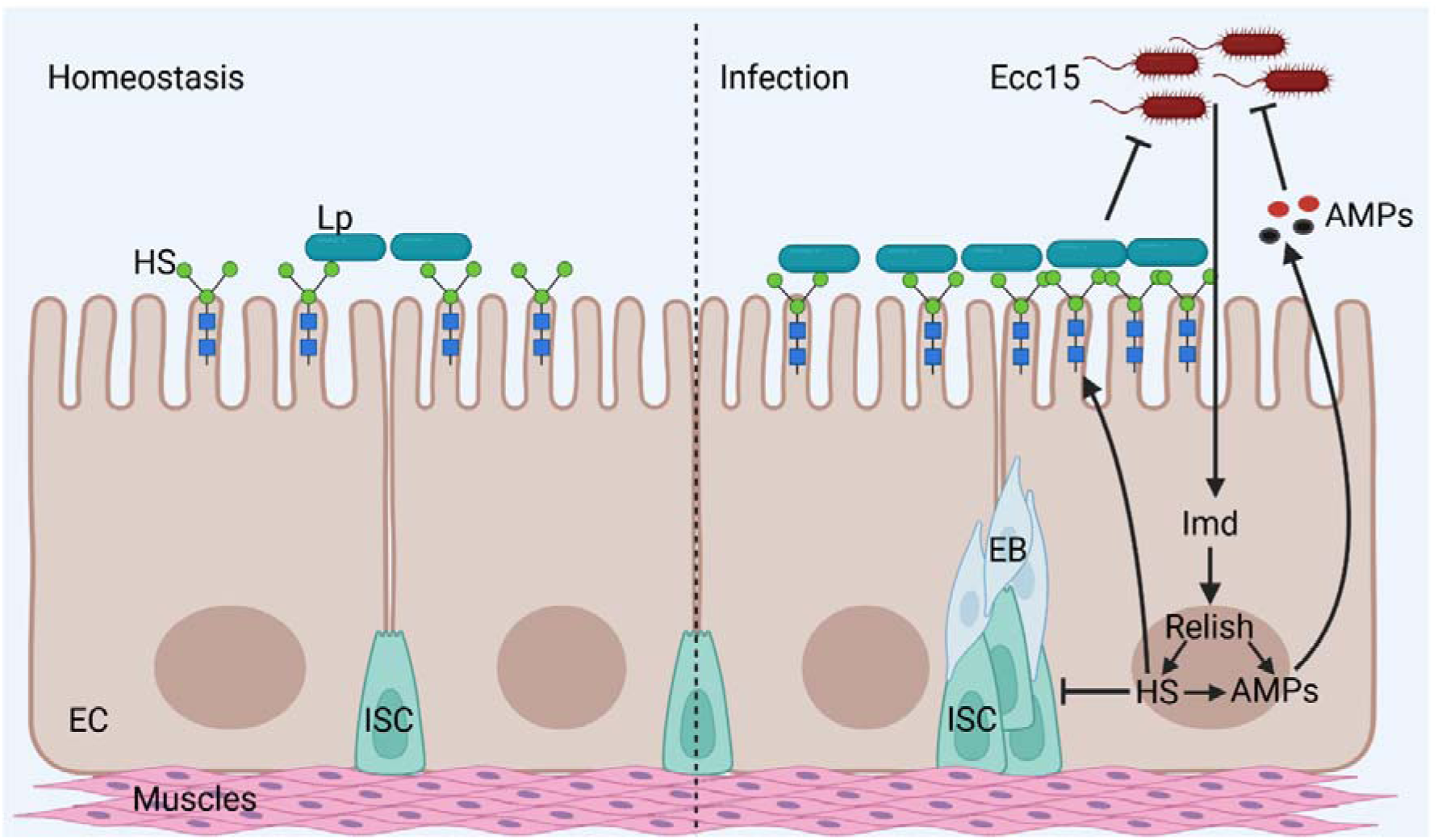
Graphical model summarizing the role of heparan sulfate in intestinal homeostasis. See Discussion for details. EC- enterocyte; EB-enteroblast; ISC-intestinal stem cell; HS-heparan sulfate, Lp-*L. plantarum*; AMPs-antimicrobial peptides; Imd-immune deficiency pathway. Created in BioRender. Iatsenko, I. (2025) https://BioRender.com/qc4xpyy

Identification of host control mechanisms of the microbiota abundance is important for prevention of negative consequences of microbiota overgrowth. We used GWAS to identify host factors that control the levels of a prominent *Drosophila* gut commensal *Lp* under basal and infection conditions. Consistent with previous studies ^24,25^, we demonstrated the importance of host genetic factors as determinants of the microbiota abundance. Considering the number of significant variants and their functional diversity, the host control of microbiota is a multifactorial process. Interestingly, we found similar GO categories enriched in *Ecc15* and sucrose gene sets suggesting that *Lp* levels are at least partially governed by the same genetic mechanisms. Enrichment of GO categories related to cell growth, adhesion, and differentiation illustrates that variation in commensal bacterial level is influenced by physical aspects of gut cell growth and development. As the microbiota affects key aspects of gut cellular physiology, including cell type, cell spacing, and epithelial turnover ^20^, the unique response of each host genotype may in turn reshape the microbial community. We also identified several GO terms specifically enriched either in *Ecc15* or sucrose gene sets. Specifically, “response to oxidative stress” and “stem cell proliferation” terms were found only in infection condition. Given that ROS and ISCs proliferation are induced by infection as defense responses ^35,36^, they might also affect the microbiota abundance. Although infection also induces the expression of AMPs, we did not find evidence for the role of immune response in the control of *Lp* abundance, likely because *Lp* is resistant to host AMPs ^48^. GO terms related to hemocyte differentiation were specifically enriched in sucrose gene set, indicating a possible role of hemocytes in the control of microbiota. Further experiments are needed to test this possibility.

Using gene knockdown by RNAi in the gut, we validated that many of the GWAS-identified candidate genes involved in various processes indeed contribute to the control of *Lp* load. We focused in detail on the role of heparan sulfate and N-acetylglucosamine metabolism in the host colonization by *Lp*. Although some *Lactobacillus* species can degrade host GAGs including heparan sulfate, and use them as nutrients ^61^, we did not find evidence for this for *Lp* strains that we used. Instead, *Lp* uses heparan sulfate for adhesion and biofilm formation. This is likely the mechanism that explains how oral administration of heparan sulfate increases *Lactobacillus* populations in rats ^62^, given the known role of GAGs in mediating adherence between *Lactobacillus* and mammalian cells ^63^.

Our work together with the other studies on the role of GAGs in host-microbiota interactions suggests that GAG availability/composition might determine the host specificity, diversity, and abundance of gut microbiome.

To further validate the role of adhesion to heparan sulfate in gut colonization and identify bacterial determinants of adhesion, we performed genetic screen for *Lp* mutants with reduced ability to bind to heparan sulfate. Importantly, 4 out of 5 genes that we hit in our screen have been previously linked to biofilm formation. For instance, extracellular transglycosylase is an aggregation promoting factor which binds to porcine mucin III via serine/threonine domain and is involved in bacterial autoCaggregation ^64^. MarR family transcription regulators control biofilm formation via regulation of gene clusters encoding putative adhesins ^65,66^. Redox- sensing transcription repressor Rex has also been implicated in biofilm formation, although the mechanism is less understood ^67^. The biofilm formation ability of bacteria is known to be affected by prophages via multiple mechanisms. Prophage P1 protein 60 or holin that we identified could facilitate the release of extracellular DNA needed for biofilm ^68^. Finally, transposon intergenic insertion between XynC (acetyl xylosidase) and xpkA (xylulose-5-P phosphoketolase / fructose-6-P phosphoketolase) was identified in our screen. Since none of the two genes have obvious link to biofilm formation, further work is needed to understand how exactly they are involved in bacterial adhesion to heparan sulfate. Notably, while all five mutants from our screen exhibited lower adhesion to heparan sulfate in vitro, they colonized flies to the similar degree as wild-type *Lp* under basal conditions but failed to expand during infection. Hence, they could not benefit from the increased availability of heparan sulfate during infection.

An outstanding question is how long infection-induced *Lp* expansion lasts and whether there are negative consequences for the host in the long-term. Since high heparan sulfate levels were associated with decreased lifespan due to oxidative stress-induced dysplasia ^43^, long-term carriage of elevated *Lp* numbers in flies recovered from infection could have detrimental effect on intestinal homeostasis. On the other hand, long-term persistence of elevated *Lp* population in the gut could provide better protection against subsequent infections. These possibilities remain to be tested.

Since many pathogens use heparan sulfate for adhesion and entry into host tissues ^52^, heparan sulfate increase induced by infection might increase host colonization by the pathogens. For example, group B *Streptococcus* (GBS) requires binding to *Drosophila* GAG to establish infection. The knockdown of GAG synthesis led to longer host survival and a lower bacterial burden ^69^. Similar GBS-GAG interactions are necessary for pathogen entry into fly nervous system ^70^. Hence, while heparan sulfate is an important mediator of gut colonization by the protective commensal, certain pathogens can exploit heparan sulfate to establish infection.

Besides the effect of heparan sulfate on microbiota, we uncovered previously unreported role of heparan sulfate in intestinal defense via the regulation of AMPs expression and intestinal repair processes. Consistent with previous studies ^56,57^, we demonstrated that disruption of heparan sulfate synthesis pathway results in intestinal hyperplasia. Such disruption of intestinal homeostasis was linked with increased susceptibility of flies to *P. entomophila* infection ^58^. Given that heparan sulfate regulates multiple signaling pathways, like Wingless (Wg), Hedgehog (Hh), epidermal growth factor receptor (EGFR), Jak-Stat, and Decapentaplegic (Dpp) with a known role in ISC proliferation ^71–74^, the role of heparan sulfate in maintaining intestinal homeostasis during infection is not surprising. The contribution of heparan sulfate to the control of AMP expression is, however, less clear. While the immunomodulatory role of heparan sulfate has been reported in mammalian cells and in *Drosophila* ^75,76^, mechanistic understanding of this process remains incomplete. One possibility could be that heparan sulfate degradation products are detected by the *Drosophila* surveillance receptors as a sign of infection, similar to how TLR4 detects heparan sulfate products in mammals ^77^. However, since *Ecc15* infection did not change overall composition of GAGs in flies, this scenario is unlikely. An alternative possibility could be that heparan sulfate modulates the function of cytokines. Given that heparan sulfate interacts with a number of endogenous secreted signaling molecules in *Drosophila* ^71^, some of these molecules might play a role in the regulation of immune response. Further understanding of signaling pathways regulated by heparan sulfate during host responses to pathogens may lead to the development of new approaches for enhancing host defenses and survival.

Overall, our study not only identified genetic determinants of microbiota load that warrant further investigation but also mechanistically dissected the role of heparan sulfate in intestinal defense and maintenance of commensal populations. These findings open possibilities for the use of heparan sulfate as an effective modulator of microbiota and host health.

## Materials and methods

### *Drosophila* Stocks and Rearing

DGRP lines were obtained from the Bloomington Stock Center. The following *Drosophila* stocks used in this study were described previously: DrosDel *w^11^*^18^ iso; *Relish^E^*^20^ iso; *w;Myo1A-Gal4, tub-Gal80TS, UAS-GFP*; *w;esg-Gal4, tub-Gal80TS, UAS-GFP*; *prospero- GAL4*; mCherry RNAi ^32,48^. The following stocks were obtained from the Bloomington Drosophila Stock Center: sdc-RNAi (51723), dally-RNAi (33952), dlp-RNAi (34089), trol-RNAi (42783), ttv-RNAi (78207), sotv-RNAi (52883), botv-RNAi (61257), slf-RNAi (34601), Hsepi-RNAi (67422), Hs3st-RNAi (28618), Sulf-RNAi (60057), ttv-RNAi (51480), Syt4 -RNAi (26730), Egfr -RNAi (25781), caps -RNAi (28020), luna-RNAi (27084), Gr58a - RNAi (64897), GstS1-RNAi (28885), nAChRalpha6-RNAi (25835), ths-RNAi (40049), Cad87A-RNAi (28716), Liprin-alpha-RNAi (53868), DIP-theta-RNAi (28654), Ptp99A- RNAi (25840), scrib-RNAi (29552), jing-RNAi (27024), Usp10 -RNAi (84055), cnc -RNAi (25984), kirre -RNAi (64918), α-Mannosidase class I a-RNAi (64994), robo3 -RNAi (29398), Fas2 -RNAi (28990), vn-RNAi (56950), DIP-epsilon-RNAi (38936), sdt-RNAi (33909), Glut1-RNAi (28645), Pde1c-RNAi (28728), CG11373-RNAi (58294), Prosap-RNAi (27284), beat-Ib-RNAi (55938), MsR1-RNAi (27529), RtGEF-RNAi (32947), toe-RNAi (29345), CG11373 -RNAi (58294), nkd -RNAi (67788), stan -RNAi (26022), CG12025 -RNAi (83936), CG12374 -RNAi (74240), rdx -RNAi (82972). Full information is stated in Table S3.

The stocks were routinely maintained at 25 °C with 12/12 h dark/light cycles on a standard cornmeal-agar medium: 3.72 g agar, 35.28 g cornmeal, 35.28 g inactivated dried yeast, 16 mL of a 10% solution of methylparaben in 85% ethanol, 36 mL fruit juice, and 2.9 mL of 99% propionic acid for 600 mL. Food for germ-free flies was supplemented with ampicillin (50 µg/mL), kanamycin (50 µg /mL), tetracycline (10 µg/mL), and erythromycin (10 µg /mL). Fresh food was prepared weekly to avoid desiccation. Female flies were used for all experiments unless stated otherwise.

### Generation of germ-free flies

16 h old embryos from each fly line were collected on grape juice plates. Embryos were rinsed in phosphate-buffered saline (PBS) and transferred to 1.5 ml tubes. All following steps were performed in a sterile hood. Embryos were placed in a 3% solution of sodium hypochlorite for 10 min. The bleach solution was discarded and embryos were rinsed three times in sterile PBS. Embryos were transferred by pipette to tubes with antibiotics- supplemented food in a small amount of 100% ethanol and maintained at 25 °C. Subsequent generations were maintained in parallel to their conventionally reared counterparts by transferring adults to new tubes with antibiotics-supplemented food. The axenic state of flies was routinely assessed by culturing.

### Colonization and infection

Ten days old female flies were starved for 2h prior to the colonization with either *L. plantarum* wild-type or with *L. plantarum* transposon mutants. Overnight bacterial cultures were adjusted to OD600=50 and mixed 1:1 with 5% sucrose. 150 μl of the mixture was placed on paper filter disks covering fly food. Flies were kept in these vials for 24h at 25°C. After 24h of colonization, flies were starved for 2h at 29°C. After starvation, flies were infected with 150 μl of either *Ecc15* or *P. entomophila* OD=200 mixed 1:1 with 5% sucrose; control flies were treated only with 2.5% sucrose. Samples were collected at desired time points.

### RT-qPCR

Gene expression was performed using 20 guts per sample. Total RNA was isolated using TRIzol reagent according to the manufacturer’s protocol. After quantification with the NanoDrop ND-1000 spectrophotometer, 500 ng of total RNA was used to perform cDNA synthesis using PrimeScript RT (TAKARA) and random hexamers. qPCR was performed on LightCycler 480 (Roche) in 384-well plates using SYBR Select Master Mix from Applied Biosystems. Expression values were normalized to RP49. Primer sequences are listed in Table S3.

### Bacteria load quantification

Pools of five flies were utilized for bacterial quantification. Total DNA was extracted from the whole flies, which were first surface sterilized in 95% ethanol for 1 minute, following the protocol of the Puregene kit (Qiagen). Flies were homogenized in 300 µl of cell lysis solution on ice using a pestle in 1.5 ml tubes and incubated at 65°C for 15 minutes, after which 100 µl of protein precipitation solution was added. The samples were vortexed and further incubated on ice for 5 minutes. Following centrifugation at maximum speed for 5 minutes, the supernatant was carefully transferred to a new tube, and DNA was precipitated by adding 300 µl of isopropanol. The DNA was pelleted through centrifugation at maximum speed for 5 minutes, and the pellet was washed with 300 µl of 70% ethanol. After air-drying, the DNA was resuspended in 240 µl of DNA hydration solution. This DNA was then diluted at a ratio of 1:4 for bacterial load quantification using specific primers listed in Table S3.

### Genome-wide association analysis

Log10-transformed *L. plantarum* load means of 8 repeats per line were used as a phenotype. Genome-wide association studies were performed using the *Drosophila* Genetic Reference Panel (DGRP) Freeze 2.0, a collection of fully sequenced inbred lines derived from a natural population. To determine genotype-phenotype associations, Log10-transformed *L. plantarum* loads were submitted to the dgrp2 webtool (http://dgrp2.gnets.ncsu.edu/)^18^. Statistical significance was assessed using ANOVA.

### Gene ontology analysis

Genome-wide top associations were used to perform gene ontology analysis. Gene Ontology Analysis: Go term enrichment for gene group lists was conducted using FlyMine ^51^. Results were filtered using a corrected p-value of <0.05 (Bonferroni).

### Heparan sulfate quantification in the *Drosophila* gut samples

20 guts were homogenized in 100 µl of 1X PBS. After dilution 1:4, 5 µl of each sample in triplicates were pipetted in the wells of 96 well plates. 180 µl of Heparin Red Ultra reagent (AMBIOS; AMS.HRU001) were added to each sample. After shaking in the plate reader for 1 min at max speed, the fluorescence was measured (Excitation at 570 nm, fluorescence emission at 605 nm) using Infinite 200 Pro plate reader (Tecan). Heparan sulfate levels were quantified by interpolating sample absorbance values onto a standard curve prepared with known concentrations of heparan sulfate.

### Heparan sulfate staining in *Drosophila* guts

To detect heparan sulfate in *Drosophila* guts, *Drosophila* guts were dissected in 1X PBS, followed by fixation with 4% paraformaldehyde (PFA) in PBS for 15 minutes at room temperature (RT). Guts were rinsed three times with 1X PBST, then nonspecific binding was blocked using 5% BSA in PBST for 1 hour at RT, and rinsed 3 times. Samples were incubated overnight at 4°C in blocking buffer with heparan sulfate antibody (AMIBIOS; F58- 10E4) at 1:100 dilution, then washed three times with 1X PBST. Next, guts were incubated for 1 hour at RT with a secondary fluorescent antibody Alexa Fluor488 (Invitrogen; 2714439) at 1:500 dilution. Nuclei were stained with DAPI (1 µg/mL in PBS) for 5 minutes. Samples were washed once with PBS, and mounted in Mowiol 4%. Images were obtained using a Leica Thunder microscope.

### GAG analysis

Germ-free flies were colonized with *L. plantarum* as described above and then subjected to different treatments: one group was fed with sucrose (uninfected control), while the other was exposed to *Ecc15* for 24h. After treatment, the flies were flash-frozen in liquid nitrogen and homogenized mechanically.

#### a) GAG isolation protocol

To begin the depilation process, each sample was weighed and placed in separate 50 mL conical tubes. Next, 20 mL of acetone was added to each tube, and the samples were gently rotated overnight to ensure thorough soaking. Following incubation, the tubes were centrifuged at 4000 g for 15 minutes at 4°C. The supernatant was carefully decanted, and the samples were then dried under a nitrogen stream to remove any residual solvent. To isolate GAGs from the protein, the samples were homogenized in 1.0 mL pronase digestion buffer (0.1 M Tris-HCl, pH 8.0, 2 mM CaCl2, 1% Triton X-100). A pronase enzyme solution was then added to each sample at a final concentration of 0.8 mg/ml and then, the samples were incubated at 55°C for 48 hours. After digestion, the enzyme was inactivated by heating for 15 minutes in boiling water. To degrade nucleic acids such as DNA and RNA, benzonase enzyme was added at a final concentration of 25 mU/mL, followed by a 2-hour incubation at 37°C. Then, the enzyme was inactivated by heating for 15 minutes in boiling water. The samples were then centrifuged at 12,000g for 15 minutes to separate undigested tissue, and the supernatant (almost 1 mL) of each sample was loaded onto a DEAE-Sepharose-2 mL packed bed column (anion exchange column) for GAG purification. The column was washed with ∼10 column volumes (20.0 mL) of loading buffer (∼pH 8 Tris Buffer, 0.1 M NaCl). The sample was applied to the column, the flow-through was reapplied, and then the column was washed with loading buffer. The sample was then eluted with 3 column volumes (6.0 mL) of elution buffer (pH 8 Tris Buffer, 2 M NaCl). Then, the eluent was dried. In order to remove excess salts and small molecular weight impurities that can interfere or suppress intensity in mass spectrometry, a desalting step was performed using a PD-10 column. The column was first equilibrated with 30 mL of water and then the sample (after dilution in 2.0 mL of water) was loaded onto the column and eluted with 3.5 mL of water. The eluate was then freeze-dried.

### b) GAG Lyase Digestion

Each sample was initially reconstituted in 200 μL of purified water, then split equally into two 1.5 mL conical polypropylene vials (100 μL each) for HS and CS digestion. For CS digestion, 3 μL of chondroitinase ABC was added to one vial, while for HS digestion, 1 μL each of heparinase I, II, and III enzymes was added to the second vial. To ensure consistency, 10 μL of the provided enzyme digestion buffer was added to each vial. The reaction mixtures were incubated for 24 hours, and upon completion, the enzymes were inactivated by heating the samples to 100°C for 5 minutes.

#### c) Aniline labelling process

The isolated material was dried in a 1.5 mL centrifuge tube. Next, 17 μL of liquid aniline (Sigma 242284) was added, followed by 17 μL of a reductant solution (prepared with 15 mg NaCNBH3 in 200 μL of a 30:70 mixture of glacial acetic acid and DMSO). The sample was pulse-centrifuged and left to react overnight at 37 °C (16 h). Afterward, it was dried in a SpeedVac for 20 hours. The reaction was resuspended in 40 μL of water, and a 10 μL aliquot was mixed with 2 μL of a 13C-aniline-tagged internal standard (10 nM) before being analyzed by HILIC-FLD-Q-TOF-MS.

#### D) HILIC-FLD-Q-TOF-MS

The HILIC method was developed on a Waters Acquity UPLC system (Waters Corporation, Milford, MA) equipped with a binary solvent manager, sample manager, and column manager. Separation was achieved on a Waters Acquity UPLC Amide BEH column (2.1 x 150 mm, 1.7 μm) maintained at 40 °C. A gradient from 91% acetonitrile to 70% acetonitrile in 50 mM ammonium formate was used to elute the GAG disaccharides. The flow rate was 0.5 mL/min, and injection volume was 2 μL. The UHPLC system was coupled to a Waters Synapt XS Q-TOF mass spectrometer equipped with an electrospray ionization source operated in negative ion mode. The following source parameters were used: source temperature 80°C, desolvation temperature 250°C, cone gas flow 50 L/hr, desolvation gas flow 1000 L/hr, capillary voltage 2.0 kV, sampling cone 35 V, source offset 4.0 V. Data were acquired in resolution mode from 200-1000 m/z.

MassLynx software (Waters Corporation) was used for molecular feature extraction and data processing. All data were acquired in the negative mode.

The molecular ion [M-H]- of each 12C-tagged aniline CS disaccharide was used to generate extracted ion chromatograms (XIC) at specific m/z values. (455.277 (D0a0- 12C tagged), 535.191 (D2a0- 12C tagged, D0a6- 12C tagged, D0a4- 12C tagged,), 615.155 (D2a6- 12C tagged, D2a4- 12C tagged, D0a10- 12C tagged). For the internal standard, the [M-H]C ion of 13C-tagged aniline CS disaccharides was used, with XIC at m/z 461.186 (D0a0- 13C tagged), 541.141 (D2a0- 13C tagged D0a6- 13C tagged, D0a4- 13C tagged), and 621.095 (D2a6- 13C tagged, D2a4- 13C tagged, D0a10- 13C tagged).

The molecular ion [M-H]- of each 12C-tagged aniline HS disaccharide was used to generate extracted ion chromatogram (XIC) at specific m/z values for visualization. (455.1579 (D0A0- 12C tagged), 535.1414 (D2A0- 12C tagged, D0A6- 12C tagged), 615.078 (D2A6- 12C tagged), 493.127 (D0S0- 12C tagged), 573.072 (D2S0- 12C tagged, D0S6-12C tagged), and 653.021 (D2S6-12C tagged)). For the internal standard, the [M-H]C ion of 13C-tagged aniline HS disaccharides was used, with XIC at 461.1779 (D0A0- 13C tagged), 541.1214 (D2A0- 13C tagged, D0A6- 13C tagged), 621.098 (D2A6- 13C tagged), 499.092 (D0S0- 13C tagged), 579.092 (D2S0- 13C tagged, D0S6-13C tagged), and 659.041 (D2S6-13C tagged).

To quantify the concentration of each disaccharide, the mass spectrum intensities were measured for all potential ion forms, including the deprotonated ion ([M-H]C), as well as the sodium-adduct ions ([M-H+Na]C [M-H+2Na]Cand [M-H+3Na]C) ratioed against 13C. These different ion forms were considered to ensure an accurate representation of the disaccharide concentrations across the spectrum. GAG analysis was performed by Creative Proteomics (USA).

### Pre-feeding flies with heparan sulfate

Germ-free flies were pre-fed either with 2 mg/ml heparan sulfate sodium (Sigma Aldrich; H7640) mixed 1:1 with 5% sucrose or with 2.5% sucrose at 29°C for 24h and then mono-colonized with *L. plantarum* using OD600 = 50 mixed 1:1 with 5 % sucrose for 24 h at 29°C. Afterward, flies were treated with 2.5 % sucrose or *Ecc15* at OD600 = 200, 1:1 with 5 % sucrose. For all the mixtures, 150 μl were placed onto paper filter disks covering the fly food surface.

### Lectin treatment

2 mg/ml of Lectin wheat (Sigma Aldrich; L9640) was fed in 1:1 mix with 5% sucrose for 24h at 29°C.

### Heparan sulfate culturing

Overnight *L. plantarum* culture was diluted to OD600 = 0.05 with either MRS or MRS supplemented with 1 mg/ml of Heparan Sulfate Sodium (Sigma Aldrich; H7640) and grown overnight. These cultures were used for colonizing germ-free flies at OD600 = 50 mixed 1:1 with 5 % sucrose for 24h at 29°C. Afterward, flies were treated with 2.5 % sucrose or *Ecc15* at OD600 = 200 mixed 1:1 with 5 %sucrose. For all the mixtures, 150 μl were placed onto paper filter disks covering the fly food surface.

### Bacterial growth

Overnight bacterial cultures were adjusted to OD600 = 0.1 and 50 μL of culture were pipetted into flat-bottom 96-well plates prefilled with varying dilutions of heparan sulfate sodium (Sigma Aldrich; H7640). Bacterial growth was measured at OD600 in an Infinite 200 Pro plate reader (Tecan) for 24h every 20 minutes.

### Binding to GAGs and biofilm formation

Overnight bacterial cultures were adjusted to OD600 = 0.1, and 50 μl of culture were pipetted into round-bottom 96-welll plates prefilled with varying dilutions of Heparan Sulfate Sodium (Sigma Aldrich; H7640) in MRS, or 100 μl of culture into pre-coated 96-round-well plates with 100 μl of varying dilutions of Heparan Sulfate Sodium in PBS for 24h prior to utilization at 4°C. Plates were incubated overnight at 37°C for *L. plantarum* or 30°C for *Ecc15* under stationary conditions.

The supernatant was discarded by inverting the plate and washing it three times in a bucket with 1X PBS. 200 µl of 0.1% solution of crystal violet were added, and the plates were incubated at room temperature for 15 minutes, ensuring consistent incubation time across all wells. Crystal violet solution was discarded by plate inversion and the plate was washed three times with 1X PBS. Plates were dried overnight, upside-down on a paper towel. 200 µl of 30% (v/v) acetic acid were added to each well to solubilize the bound crystal violet and incubated at room temperature for 15 minutes while gently agitating with the Belly Dancer at a set rpm. Finally, 100 µl of the solubilized dye solution from each well were transferred to a new flat bottom 96-well plate to measure the absorbance at 570 nm using a microplate reader; all reads were performed in an Infinite 200 Pro plate reader (Tecan).

### Biofilms screening

96-well round plates were coated with 100 μl of 2 mg/mL of heparan sulfate sodium in 1X PBS for 24h prior to utilization at 4°C. *L. plantarum* cultures of the previously generated *Lp* library ^48^ were recovered by incubating them overnight at 37°C under stationary conditions. Overnight bacterial cultures were diluted 1:100 in MRS supplemented with 5 mg/ml of Erythromycin in the heparan sulfate-coated plate, and the initial OD was recorded at OD600. Plates were incubated overnight at 37 °C, and the final OD 600 was read. Biofilm formation was recorded as stated above.

To identify samples with normal growth but low biofilm formation, we filtered the dataset to include only those with optical density values within ±10% of the mean growth (mean_growth) while retaining samples with crystal violet (CV) staining values below 80% of the mean biofilm formation (mean_biofilm). The following formula was used for the filtering criteria: 0.9 x mean_growth ≤ OD ≤ 1.1 x mean_growth and CV < 0.8 x mean_biofilm

### Bacterial growth in modified MRS media

To assess the ability of *Lp* to utilize heparan sulfate as a carbohydrate source, we used the following modified MRS media adapted from ^78^: 10 g L^−1^ tryptone, 10 g L^−1^ beef extract, 5 g L^−1^ yeast extract, 2 g L^−1^ tri-ammonium citrate, 3 g L^−1^ sodium acetate, 0.1 g L^−1^ magnesium sulphate heptahydrate, 0.038 g L^−1^ manganese sulphate monohydrate, 2 g L^−1^ dipotassium phosphate, 1 mL^−1^ Tween 80, pH 6.2. Control medium contained 20 g L^−1^ of glucose, while experimental media were supplemented with 20 g L^−1^ of heparan sulfate or 20 g L^−1^ of N- acetyl glucosamine. Overnight *L. plantarum* culture was grown overnight in standard MRS broth. Next day, the culture was diluted to OD 0.01 with each of the modified MRS media to be tested. Adjusted *L. plantarum* cultures (180 µl) were pipetted in the wells of 96 well plate (at least 3 wells per strain/treatment) and their growth was assessed by measuring OD600 in the plate reader at 37 °C.

To test whether *Ecc15* can increase the usability of heparan sulfate by *L. plantarum*, we pre-treated heparan sulfate with LB media or with *Ecc15* culture for 3 h at 30 °C. After incubation, *Ecc15* was removed via centrifugation and filtration and treated heparan sulfate solution was used to supplement modified MRS media as a sole carbohydrate source. *L. plantarum* growth was measured in these media as described above.

### Minimal Inhibitory Concentration

Overnight bacterial cultures were adjusted to OD600 = 0.1 and diluted 1:100. 50 μl of culture were pipetted into flat-bottom 96-well plates prefilled with varying dilutions of Polymyxin B.

### Heparan digestion by *Ecc15*

*Ecc15* overnight culture (100 ml) was collected by centrifugation. Pellets were suspended in 1 ml Tris/NaCl Buffer (20 mM Tris HCl, 100 mM NaCl, 0.5mM EDTA supplemented with Protease Inhibitor Cocktail). The suspension was incubated with or without Lysozyme (50 µg/ml final) for 30 min at ambient temperature. DNA was digested with 10U Benzonase-Nuclease (Merck) for a further 30 min in the presence of 2 mM MgCl2 and the suspensions were centrifuged at 10.000xg for 5 min. Pellets (“P +/-Lysozyme”) were suspended in 100 µl Heparinitase-Digestion Buffer (100 mM NaAc, 2 mM Calcium Acetate, pH 7.5). Supernatants (“S/N +/-Lysozyme”) were diluted fivefold in digestion buffer and concentrated via Ultrafiltration (Ultrafree 10K, Millipore) to a final volume of 100µl.

### Heparan digestion

20 µl of the lysed samples (supernatant and pellet) were incubated with 1 µg (1µl) of Heparan sulfate at 32°C for 2 hours. The reaction mixtures were stopped by adding 200 µl chloroform/methanol (2:1, v/v) and 100 µl of water. After vertexing and centrifugation, the aqueous upper phase was removed and dried under vacuum in a SpeedVac- Centrifuge.

### Derivatization with 2-Aminocaridone (AMAC) ^79^

For the AMAC labeling, the dried samples were mixed with 20 μL of 0.1 M AMAC solution in DMSO/glacial acetic acid (17:3, v/v) and incubated at room temperature for 15 min. To the reaction mixture, 20 μL of 1 M aqueous sodium cyanoborohydride was added. The reaction mixture was incubated at 45 °C for an additional 2 h. The reaction was centrifuged to obtain the supernatant for UPLC analysis.

### UPLC

Reversed-phase UPLC was performed using an Acquity UPLC-System from Waters, fitted with a HSS-T3 column (2.1 x 100mm, 1.8µm). A linear gradient from buffer A (10 mM Ammonium Acetate pH 5.6 bicarbonate) to 50% acetonitrile in Buffer A over 7min at 45°C and a flow rate of 0.4 ml/min was applied. Eluted compounds were detected by fluorescence (ex 425 nm, em 532 nm) and by ESI-MS detection (waters QDa).

The QDa was operated in an electrospray negative ion mode by applying a voltage of 0.8 kV. The cone voltage was set at 15 V. The probe temperature was set at 600 C. A full mass spectrum between *m*/*z* 100 and 1200 was acquired at a sampling rate of 8.0 points/sec.

### GFP measurement

10 guts were mechanically homogenized in 200 µl of Schneider’s Drosophila medium (Gibco; 21720001) and incubated for 15 minutes at 25 °C. The cells were then spun down at 1200 rpm for 5 minutes and resuspended in 500 µl of 1X PBS with 2% FBS. A total of 150 µl of the cell suspension was transferred in duplicates and measured in the plate reader at an excitation wavelength of 485 nm and an emission wavelength of 520 nm.

### Imaging

*Drosophila* guts were carefully dissected in 1X PBS and then fixed using 4% paraformaldehyde (PFA) in PBS for 15 minutes at room temperature. Nuclei were stained with DAPI at a dilution of 1:500 (1 µg/mL in PBS) for 5 minutes, followed by a single wash with PBS. The samples were then mounted in Mowiol 4%. Imaging was performed using a Leica Thunder microscope.

### Statistical analysis

Unless otherwise stated, all statistical analyses were performed in R version 4.3.1. In vitro: bacterial growth experiments, MIC, GFP quantification, biofilm formation, binding quantification enclose three technical triplicates and three biological duplicates. In vivo, experiments like colonization, infection, pre-feeding assays, gene expression quantification, bacterial loads, survivals, and heparan sulfate quantification data were derived from three experiments and four biological repeats. Survival analysis was carried out using the Wilcoxon method and the Log Rank test using the R package survminer. Pairwise and two independent group comparisons were performed with a t-test; Kruskal-Wallis and Bonferroni–Holm post hoc tests were used for multiple group comparisons. Statistical significance is indicated as follows: non-significant (ns) > 0.05, *p < 0.05, **p < 0.01, ***p < 0.001, ****p < 0.0001. Error bars represent the mean’s standard deviation (SD) or error (SE) as indicated per experiment. Data visualization was performed with the R packages ggplot2, dplyr, reshape2, tidyverse, ggpubr, ggrepel, devtools, rstatix and multcomp. Graphic draws were created using Biorender and adapted into Illustrator 2025.

## Acknowledgements

We are grateful to the Bloomington Drosophila Stock Center (NIH P40OD018537) for fly stocks. This work was supported by the Max Planck Society. I.I. also acknowledges the funding from the Deutsche Forschungsgemeinschaft (grants IA 81/2-1 and IA81/3-1) and from the Boehringer Ingelheim Foundation.

## Author Contributions

AAR – investigation, formal analysis, data curation, methodology, visualization, supervision, writing – original draft preparation, writing – review and editing; MR – investigation, validation, writing – review and editing; JA – investigation, writing – review and editing; DF investigation, resources; RH – investigation, methodology, writing – review and editing; II conceptualization, funding acquisition, project administration, supervision, resources, writing – original draft preparation, writing – review and editing.

## Supplemental Information

Table S1. Excel table listing GWAS-identified SNPs and candidate genes associated with L. plantarum abundance under uninfected conditions.

Table S2. Excel table listing GWAS-identified SNPs and candidate genes associated with L. plantarum abundance under infection conditions.

Table S3. Excel table listing fly lines, primers, and bacterial strains used in this study.

---

**Figure S1.**
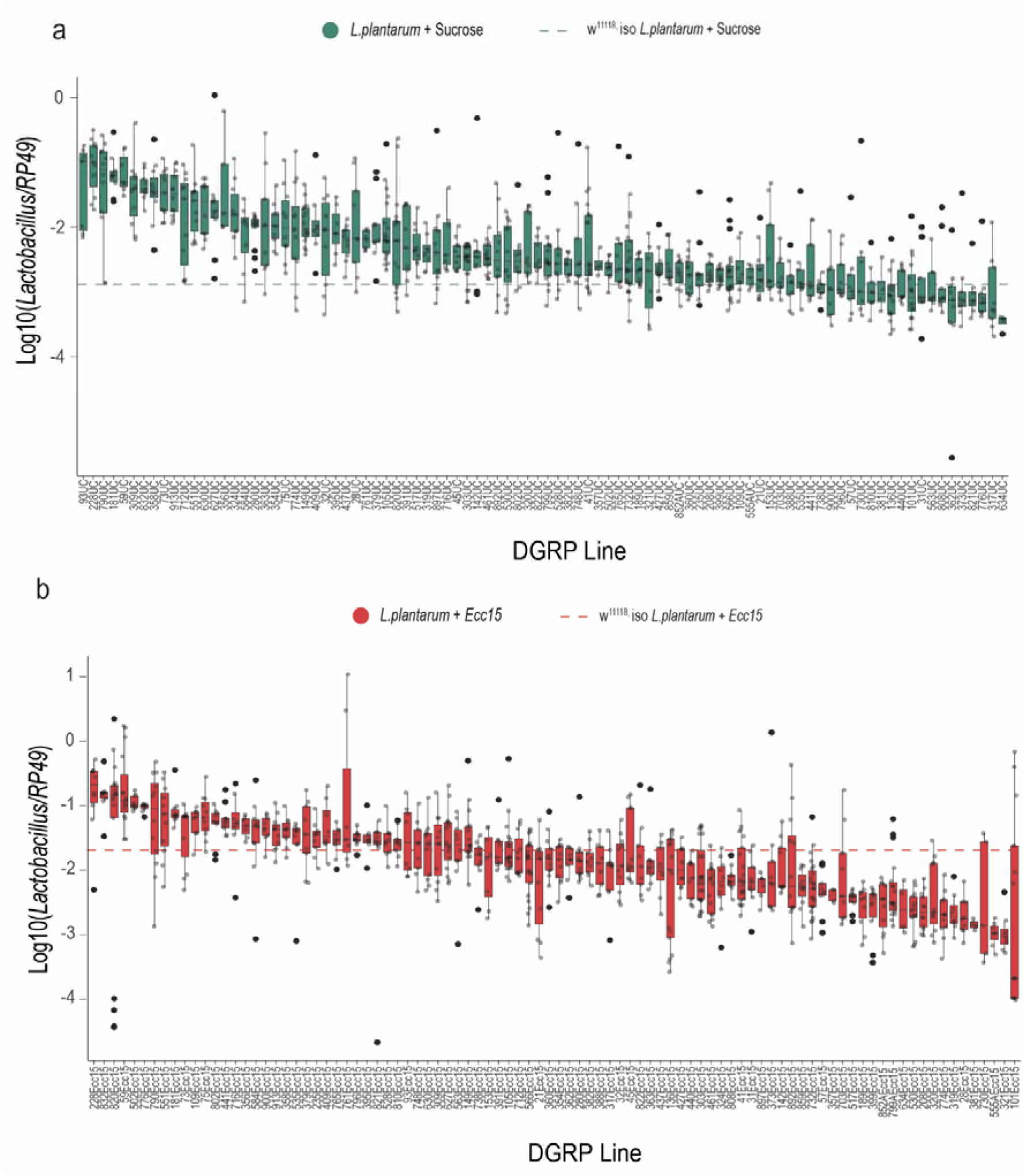
*L. plantarum* levels vary among the DGRP lines. (**a**) *L. plantarum* loads among DGRP lines colonized with *L. plantarum* and treated with 2.5% sucrose for 6h. (**b**) *L. plantarum* loads among DGRP lines colonized with *L. plantarum* and infected with *Ecc15* for 6h.

**Figure S2.**
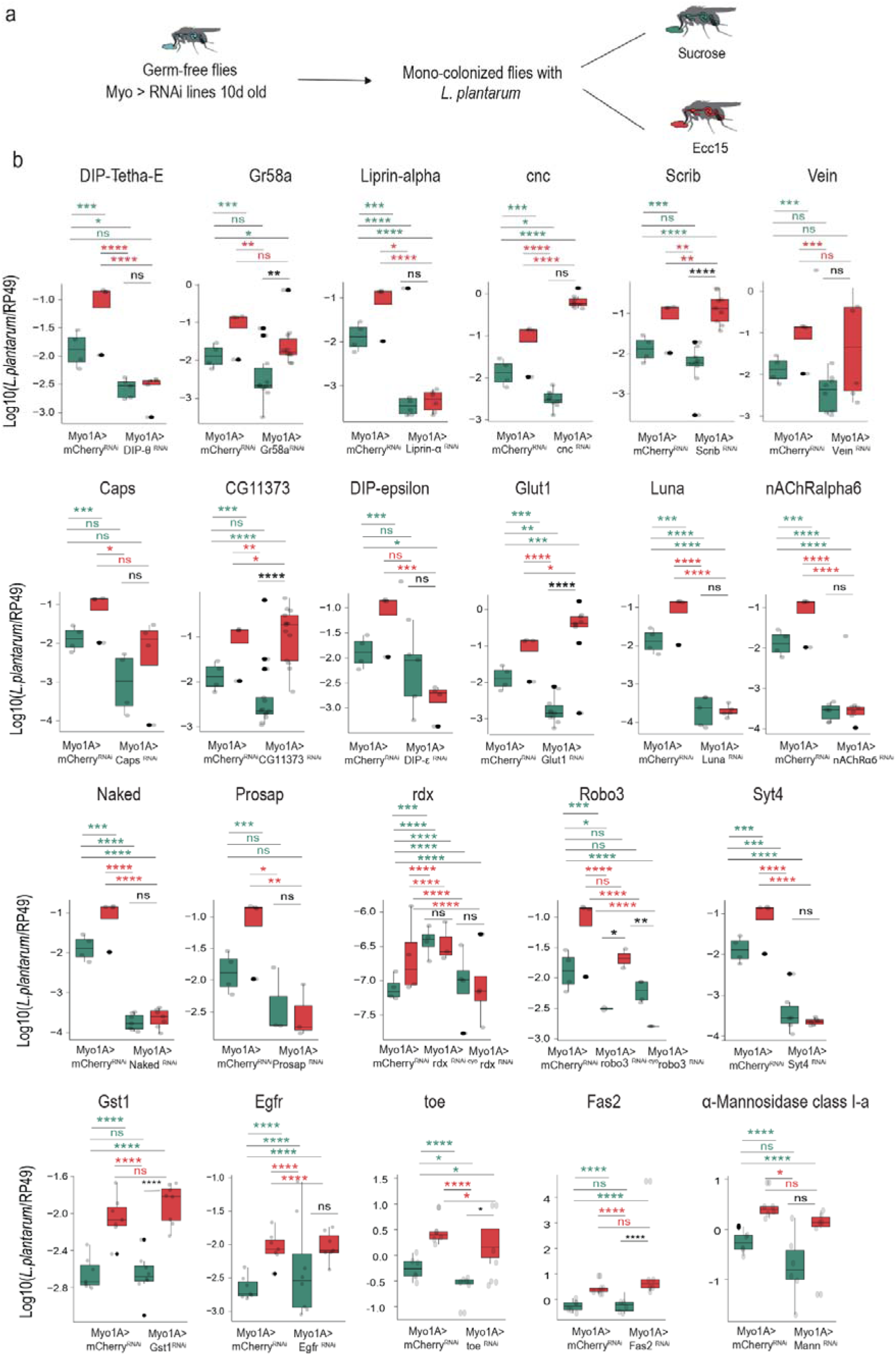
Experimental validation of genetic variants derived from the GWAS analysis. (**a**) Scheme of experimental procedure followed to validate the role GWAS-identified candidate genes in the control of *L. plantarum* abundance. (**b**) Levels of *L. plantarum* in RNAi lines that knock down the most significant GWAS-identified candidate genes in the gut.

**Figure S3.**
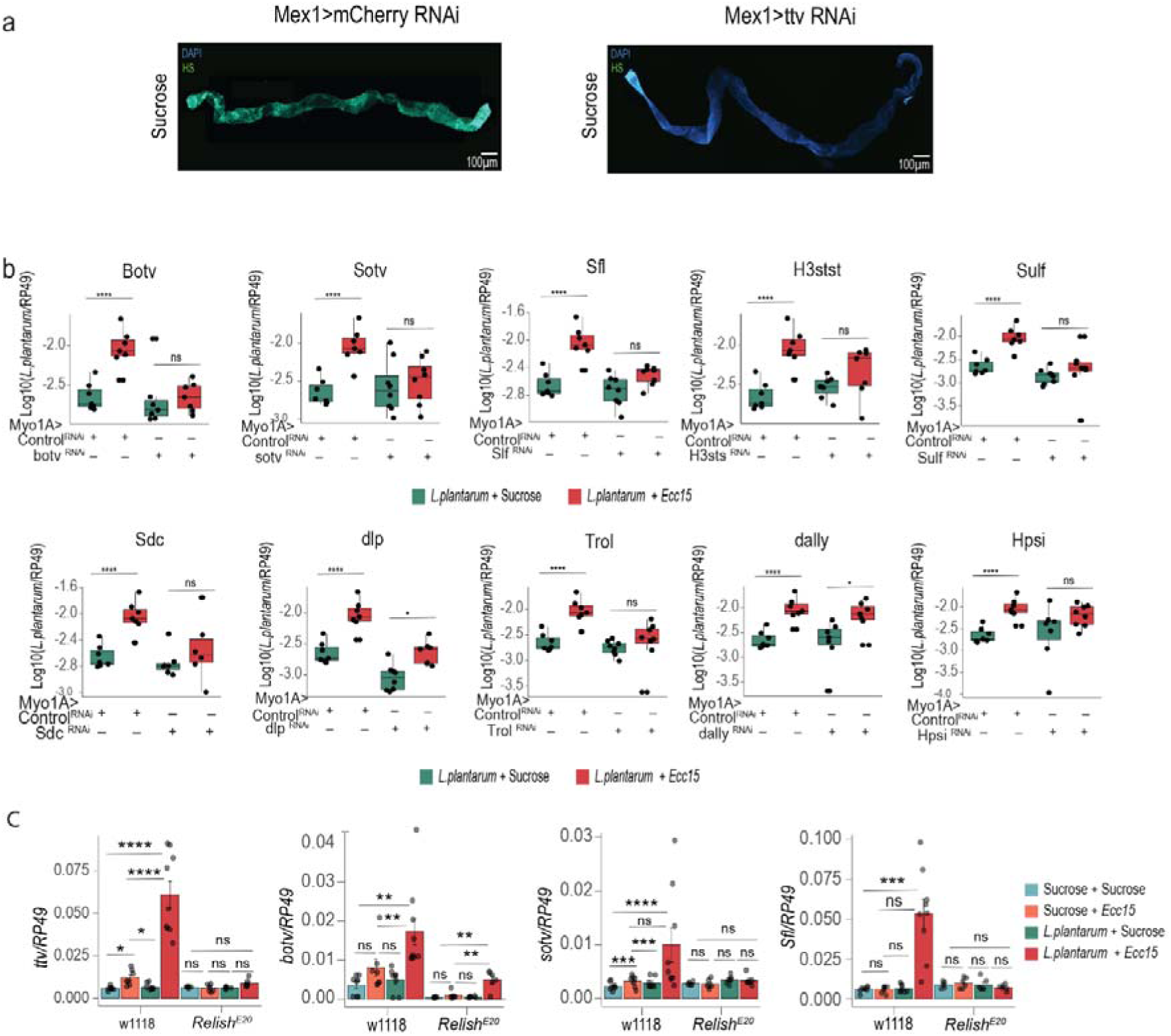
The heparan sulfate pathway mediates *L. plantarum* abundance during infection through the Imd pathway. **(a)** Representative images of heparan sulfate staining in guts from Myo1A>mCherry RNAi and Myo1A>ttv RNAi flies. (**b**) Loads of *L. plantarum* in Myo1A>attp40 RNAi (control) and in RNAi lines with knockdown of genes associated with the heparan sulfate synthesis pathway. (**c**) Expression of heparan sulfate pathway genes in wild-type and *RelishE20* germ-free flies treated with sucrose only, colonized with *L. plantarum*, and treated with sucrose again or infected with *Ecc15* and collected at 6h (n=8).

**Figure S4.**
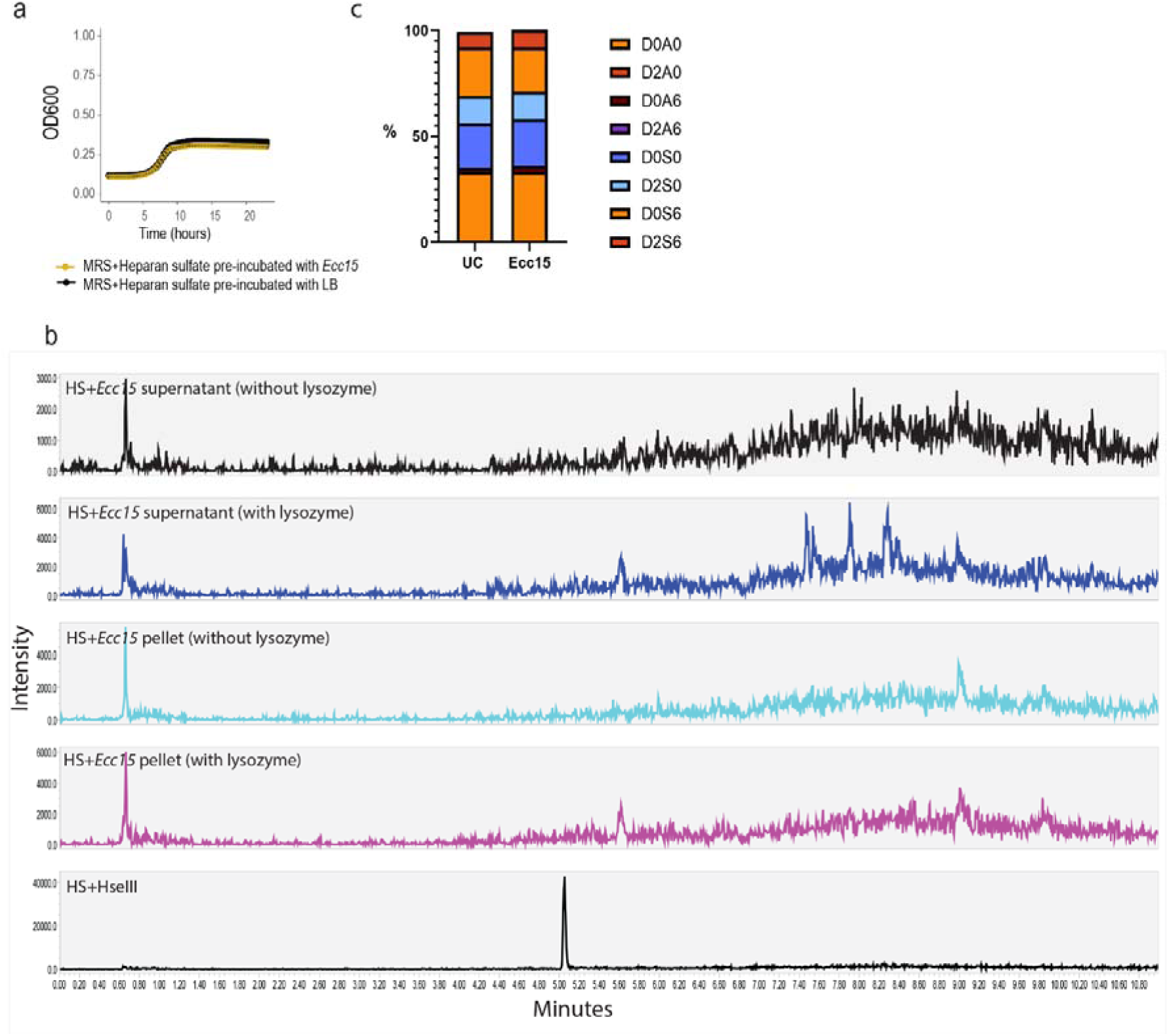
*Ecc15* does not change *Drosophila* Glycosaminoglycan composition. **(a)** *L. plantarum* growth in modified MRS medium supplemented with heparan sulfate pre-incubated with LB media or *Ecc15* bacteria. (**b**) Detection of heparan sulfate digestion products by Reversed-phase UPLC. 2-AMAC-derivatized fluorescent digestion products were detected only in a positive control sample (as illustrated by the peak) but not in *Ecc15*-treated samples. (**c**) Composition of glycosaminoglycans in uninfected and *Ecc15*-infected flies analysed by HILIC-Q-TOF-MS (see methods). Proportion of each disaccharide (in %) relative to all detected disaccharides is shown.

**Figure S5.**
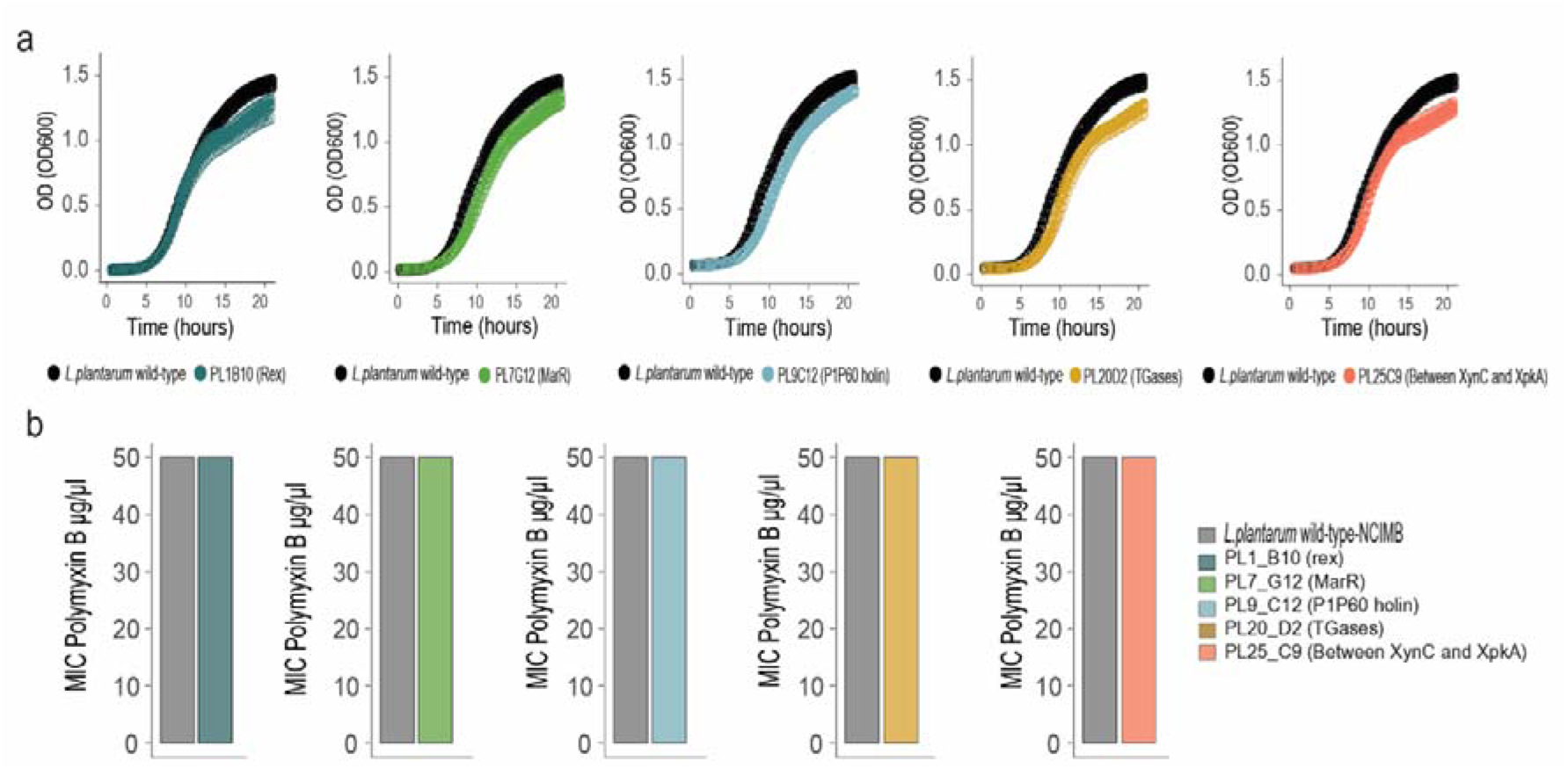
Transposon mutants showed unimpaired growth and resistance to polymyxin B. (**a**) Growth curves of *L. plantarum* wild-type and the transposon mutants with reduced biofilm formation. (**b**) Sensitivity of *L. plantarum* wild-type and the transposon mutants to polymyxin B in Minimal Inhibitory Concentration assay. The experiment was repeated three times with identical results, therefore, no error bars are shown.

**Figure S6.**
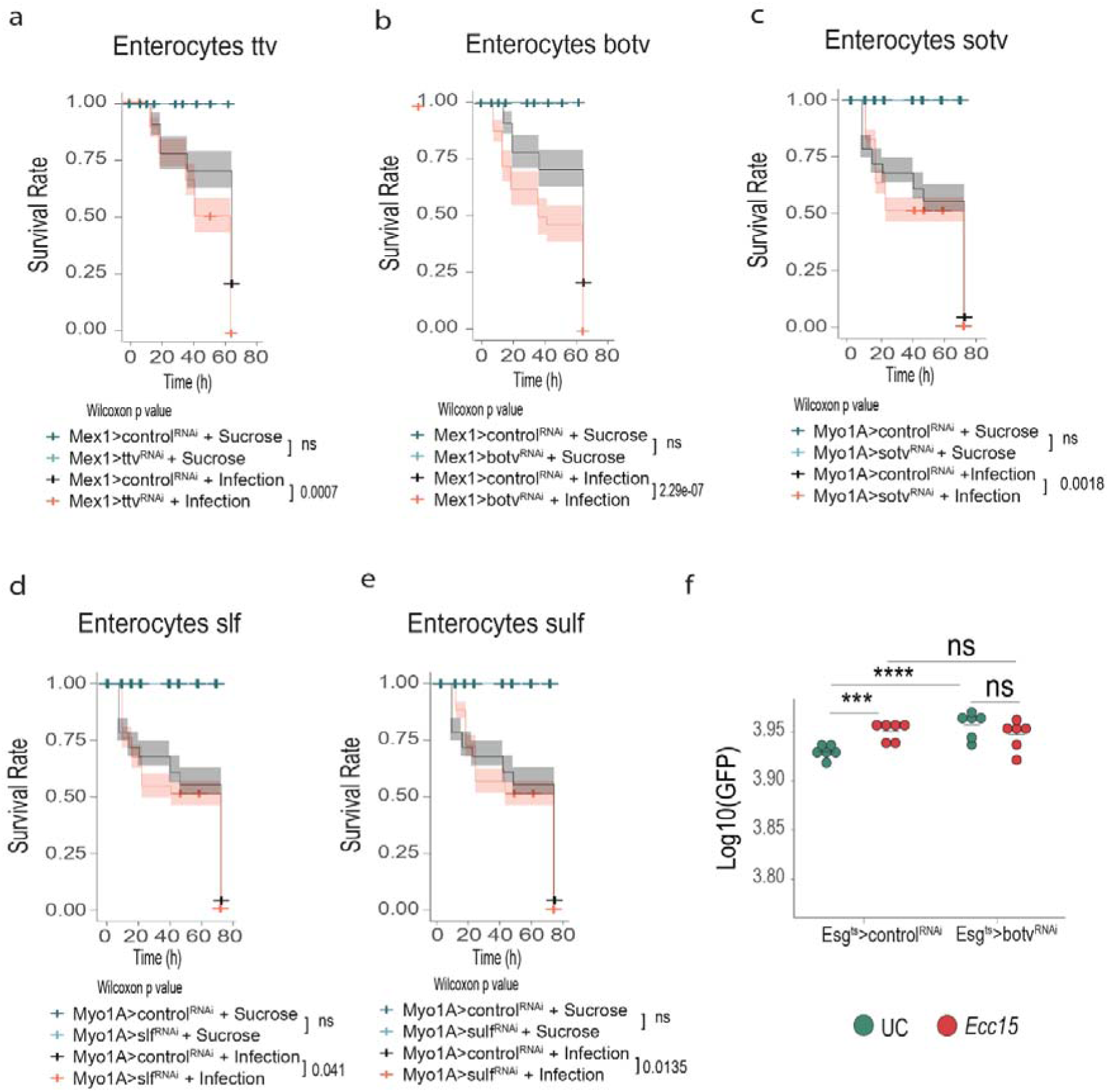
Heparan sulfate pathways contributes to intestinal defense against pathogens. **(a-e)** Survival curves of RNAi lines with enterocyte-specific knockdown of heparan sulfate pathway genes: *ttv* (a), *botv* (b), *sotv* (c), *slf* (d), *sulf* (e) exposed to either sucrose or *P. entomophila* infection. (f) Quantification of escargot positive cells (GFP positive cells) in Esg>attp40 RNAi and Esg>botv RNAi treated with either sucrose or *Ecc15* for 16h (n=6).

